# Cyclical content exchange in hydrogel-core synthetic cells

**DOI:** 10.1101/2025.07.16.665112

**Authors:** Aileen Cooney, Wancheng Zhang, Lorenzo Di Michele, Yuval Elani, Tomoaki Matsuura

## Abstract

Synthetic cells lack the complex protein machinery that living cells use to control molecular traffic across their membranes, making programmable exchange with the environment a central challenge in bottom-up synthetic biology. Here, we introduce a protein-free approach for dynamic content exchange in liposome-based synthetic cells through a membrane-hydrogel composite architecture. We assemble the synthetic cell by crosslinking a polymer hydrogel within the lumen of giant unilamellar vesicles (GUVs) and chemically tethering it to the inner membrane leaflet. The resulting mechanical coupling enables a previously unexplored response through which the membrane can be reversibly permeabilized and re-sealed upon osmotic swelling and de-swelling of the hydrogel. This mechanism of membrane remodeling enables cyclic exchange of small molecules and macromolecules, offering a versatile route for dynamic regulation of intracellular biochemistry that does not rely on reconstituted protein pores.

## Introduction

Synthetic cells are increasingly complex, micron-scale systems encapsulating biomolecular machinery and mimicking aspects of cellular functions. These systems can incorporate a diverse range of components, including synthetic and natural polymers, biomolecules, and inorganic particles.^1–5^ Synthetic cells exhibit functions that range from protein synthesis,^6–8^ molecular transport^9–11^ and motility,^12–14^ to engineering novel stimuli-responsive communication pathways.^10,15–18^ These advances can be exploited across diverse applications, including protein engineering,^19^ biocomputing,^20^ precision medicine,^21^ and origin-of-life research.^22^

Synthetic cells require the segregation of their contents from the environment, with typical compartmentalization systems including polymersomes,^23,24^ liposomes,^25,26^ proteinosomes,^27–29^ coacervates,^30,31^ condensates,^32^ water-in-oil droplets^33,34^ and hydrogel particles (microgels).^35–37^ These compartmentalized architectures enable the maintenance of intracellular reaction networks and can also mediate material exchange in response to environmental stimuli.

Controlled content exchange is an essential function of living and, therefore, synthetic cells, enabling the replenishment of chemical energy carriers, supplementation of substrates for metabolic pathways, efflux of reaction products, and modulation of cellular composition, including macromolecules. Whilst membrane proteins can support highly specific cargo transport, their use introduces dependence on complex expression systems, folding pathways, and narrow physicochemical tolerances.^38^ Hence, only a limited number of tractable membrane proteins have been validated in synthetic cells, ^39–41^ and it is difficult to precisely initiate, halt, and re-initiate cargo influx and efflux. Alternative switchable transporters based on DNA nanostructures have been demonstrated,^42–44^ but it remains challenging to mediate the transport of macromolecular cargoes. This transport bottleneck prevents the robust implementation of synthetic cell functions such as sensing, communication, collective behaviours, and the harvesting of environmental energy and substrate sources, which in turn prevents the maturation of synthetic cell technologies into a translational platform. Addressing this urgent limitation motivates the search for simpler, protein-free strategies to achieve on-demand content exchange in synthetic cells.

Here we present a synthetic cell architecture in which controlled transport emerges from the mechanical and chemical coupling between lipid membranes and an internal hydrogel. Hydrogels are hydrophilic polymer networks that have been shown to enhance synthetic cell mechanical stability^45,46^ or induce controlled changes in lumen viscosity.^47^ We sought to use the swelling and shrinking dynamics of the hydrogel^48^ to control the transport function of the synthetic cells. Changes in osmolarity act as the external stimulus, driving reversible hydrogel swelling and shrinking that dynamically open and close the membrane for the uptake and release of cargoes ranging from small molecules to proteins and DNA. Hydrogel–membrane chemical conjugation, in particular, enables membrane resealing and restoration of barrier function upon deswelling, unlocking a reversible mode of macromolecular transport that was not previously possible in synthetic cells. This provides a simple and reversible route to transport and fuel delivery without reconstructing complex biological machinery.

## Results and Discussion

### In-situ gelation of synthetic cell interiors

First, we set out to construct composite synthetic cells consisting of a GUV containing a hydrogel core in its lumen, with the aim of engineering new transport functions by exploiting the composite properties of the hydrogel interior and lipid membrane. We used the emulsion phase transfer method^49,50^ to prepare 1-palmitoylpalmitoyl-2-oleoyl-sn-glycero-3-phosphocholine (POPC) GUVs encapsulating 20 kDa, 4-arm thiolated PEG polymers ((PEG-SH)4), which were then crosslinked through an enzymatic process (Fig. 1a). In brief, glycyl-L-tyrosine and horseradish peroxidase (HRP) were co-encapsulated with (PEG-SH)_4_ in the GUV lumen to facilitate intermolecular disulfide bond formation through a phenolic radical intermediate.^51^ The gelation state of the (PEG-SH)_4_ within synthetic cells was monitored by labelling it with AlexaFluor488- or AlexaFluor647-maleimide to yield the fluorescently tagged PEG product (AF488-PEG or AF647-PEG). Unless noted, the synthetic cells were immobilised on the glass slides via biotin-streptavidin linkages (see **Methods**).

**Figure 1.**
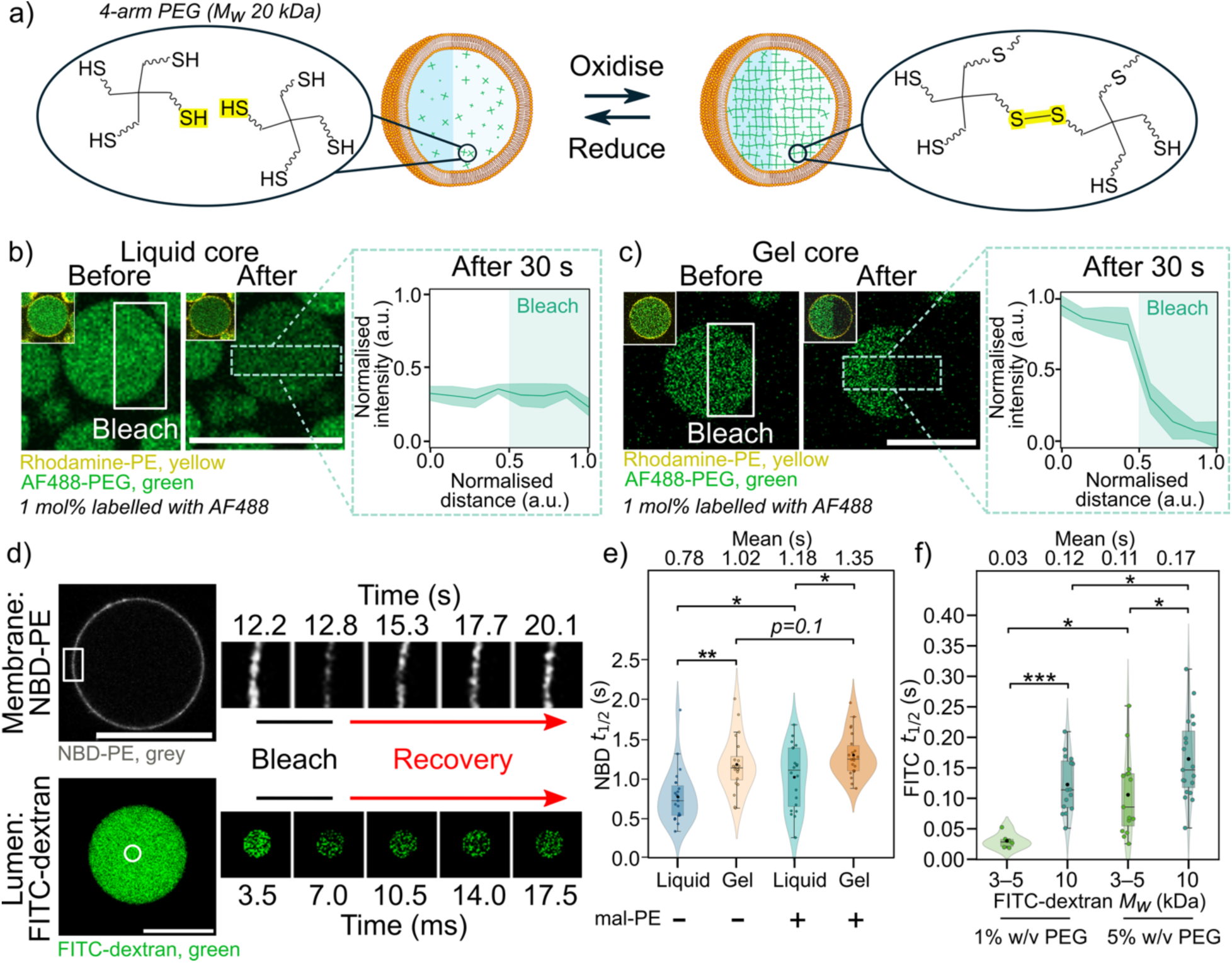
Characterisation of gel- and liquid-core synthetic cells, and the associated mobility of the membrane and the cargo in the lumen. (a) Schematic illustrating the assembly/disassembly of a gel network in a synthetic cell lumen by oxidation/reduction of encapsulated (PEG-SH)_4_. (b,c) Confocal micrograph (left) and mean plot profile (right) showing exemplar synthetic cells before and after photobleaching of the AF488-PEG in liquid-(b) and gel-core (c) synthetic cells, with the rhodamine/AF488 overlay of the same image shown in the inset. The bleach ROI is indicated by the solid box and the plot profile ROI by the dashed white box. *n*=8 (liquid), 7 (gel); lines with shaded regions in the line plots indicate mean ± 1 s.d. (standard deviation). Fluorescence intensity was normalized by subtracting the background and scaling to mean initial intensity=1. (d) Exemplar FRAP experiments showing bleaching and recovery in the ROI taken at the equatorial plane of the synthetic cell for NBD-PE in the membrane (upper) or FITC-dextran in the lumen (bottom). (e) Violin plots indicating *t*_1/2_ of NBD-PE in the membrane for liquid- and gel-core synthetic cells in the presence and absence of a mal-PE lipid anchor in the inner leaflet (*n*=20 for all). (f) Violin plots indicating *t*_1/2_ of 3-5 kDa or 10 kDa FITC-dextran in the lumen of gel-core synthetic cells with 1% or 5% w/v (PEG-SH)_4_ (*n*=8, 16, 15, 19 (l-r)). *p* > 0.05 (n.s.), *p* < 0.05 (*), *p* < 5 x 10^-3^ (**), *p* < 5.4 x 10^-4^ (***) and *p* < 5 x 10^-5^ (****). All *p*-values are provided in the SI. All scale bars=20 µm.

We used photobleaching (Figs. 1b-c) and membrane lysis (Fig. S1) experiments to determine the physical state of the synthetic cell lumen. A portion of the fluorophores within a region of interest (ROI) in the lumen was photobleached using a laser-scanning confocal microscope (LSCM). In synthetic cells without HRP and retaining a liquid lumen (“liquid-core synthetic cells”), the fluorescence intensity decreased throughout the entire lumen due to rapid diffusion of the AF488-PEG in and out of the exposed ROI (Fig. 1b, Movie S1). Conversely, in synthetic cells encapsulating HRP (“gel-core synthetic cells”), photobleaching was limited to the exposed ROI, indicating immobilization of the PEG monomer and thus confirming gelation (Fig. 1c, Movie S2). Furthermore, when the detergent Triton-X-100 was added to gel-core synthetic cells, release of encapsulated calcein and lysis of the rhodamine-labelled membrane (rhodamine-PE) was observed, while the AF647-PEG hydrogel core remained intact (Fig. 2g, Fig. S1, Movie S3). Conversely, liquid-core synthetic cells completely lysed upon breakdown of the membrane, resulting in complete content release (Fig. 2e, Fig. S1, Movie S4). These results indicate that a gel network was successfully formed in the synthetic cell lumen in the presence of HRP.

**Figure 2.**
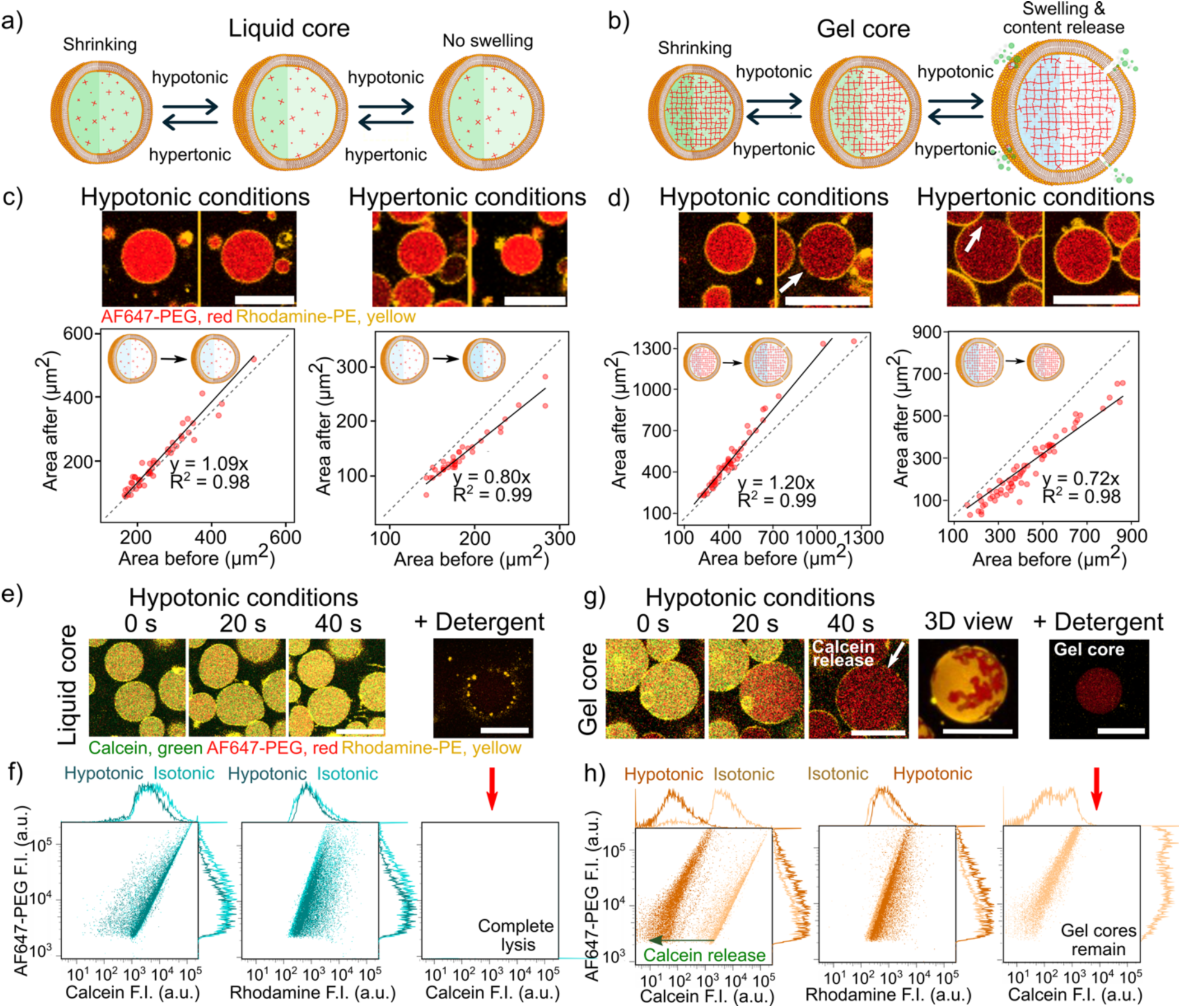
The hydrogel network within gel-core synthetic cells enhances the vesicle shrinking and swelling response to changes in osmolarity. All data indicate AF647-labelled PEG. Calcein was initially encapsulated in the lumen and rhodamine-PE was used to visualize the membrane. (a-b) Schematics indicating the effect of osmolarity changes on liquid-(a) or gel-core (b) synthetic cells. Considerable hypotonic swelling of gel-core synthetic cells introduces membrane fissures that facilitate the release of encapsulated material, shown as a green fluorophore (b, right). (c-d) Plots indicating the size increase and decrease of liquid-(c) and gel-core (d) synthetic cells in hypotonic and hypertonic buffer, with the size taken as the cross-sectional area of the synthetic cell. The dotted grey line indicates *y=x* (*i.e.* no change in size); the black line indicates a linear fit to the data points with a constraint through the origin. (*n*=49, 39 (liquid-core: left, right), *n*=66, 52 (gel-core: left, right)). (e, g) Confocal micrographs over time and (f, h) flow cytometry (FC) data indicating fluorophore intensities (F.I.) under isotonic and hypotonic (swelling) conditions for liquid-core (e, f) and gel-core (g, h) synthetic cells, as well as the response to the addition of Triton-X-100. *n*=10,000 for all FC data. (g) includes an additional confocal micrograph of the 3D reconstruction of a representative gel-core synthetic cell after exposure to hypotonic condition, with large membrane fissures visible in the membrane (yellow) leading to exposure of the gel core (red). All scale bars=20 µm.

### Gel core impacts diffusion in both the membrane and lumen

To reveal the physicochemical characteristics of the gel-core synthetic cells, we next sought to explore how the material state of the vesicle interior—liquid or gel—influences both the properties of molecular cargoes in the lumen, and those of the surrounding membrane. Fluorescence Recovery After Photobleaching (FRAP) experiments indicated that the internal gel network reduced the diffusion of both lipids in the membrane and solutes in the lumen (Figs. 1d-f).

The half maximal recovery time, *t_1/2_*, measured for NBD-labelled phosphoethanolamine (NBD-PE), was significantly larger in gel-core compared to liquid-core synthetic cells with identical lipid composition and buffer conditions (Fig. 1d-e, Movie S5), indicating that lumen gelation reduces lipid mobility. The estimated diffusion coefficients (Table S1) are in close agreement with previously reported values of NBD-cholesterol in vesicle membranes (0.53 ± 0.04 µm^2^ s^-1^)^52^ and with the typical order of magnitude of 1 μm^2^ s^-1^ for NBD-PE in PC GUVs.^53,54^ The trend is also consistent with the expectations that a solid interface in contact with a membrane reduces lipid diffusivity due to frictional coupling.^54–56^ The effect of N-(3-maleimide-1-oxopropyl)-L-α-dioleoyl phosphatidylethanolamine (mal-PE) is explored in further detail below (Figs. 4,5).

Similarly, the gel core was seen to impact the diffusion of soluble species in the lumen (Fig. 1f). FRAP showed reduced diffusion for 10 kDa compared to 3-5 kDa dextran, whilst increasing (PEG-SH)_4_ concentration reduced the mobility of both dextran species. Diffusion constants estimated from the *t_1/2_* were consistent with previous studies (Table S2).^57,58^ The mobility of the small molecule calcein (M_w_ 0.6 kDa) and the larger AF647-labelled ovalbumin (OA647) (M_w_ ∼45 kDa) was also investigated (Fig. S2, Movie S6). Calcein fluorescence decreased uniformly across the entire lumen during photobleaching, indicating its rapid diffusion. In contrast, OA647 fluorescence was completely photobleached in the exposed ROI but only reduced by 60% in the non-exposed ROI, indicating an immobile OA647 fraction of ∼40% (Fig. S2). This immobilization of OA647 was also evident in membraneless gels (Fig. S3, Movie S8) and is likely due to covalent bonding between free thiols in the gel network and cysteine residues on the protein surface.^59,60^ This ability to chemically immobilize target molecules in the lumen is desirable for pathway engineering in synthetic cells.^61,62^

The gel-core synthetic cell also features the dynamic ability to switch between gel and liquid states by exposure to membrane-permeable redox agents. This was evidenced by the dissipation of the photobleached ROI within gel-core synthetic cells upon addition of dithiothreitol (DTT) (Figs. S4, Movies S9, 10a), and subsequent gel reassembly upon addition of hydrogen peroxide (Movie S10). Membrane integrity was maintained throughout, as demonstrated by the retention of encapsulated calcein (Fig. S4d). Such switching behaviour may be valuable for dynamically altering the mobility of moieties inside the lumen, which could influence biomolecule functionality or reaction kinetics.

### Reversible swelling of gel-core synthetic cells drives the release and uptake of diverse cargo

Gelation of the lumen provides a route towards cargo transport in synthetic cells in response to osmotic changes. We exposed gel- and liquid-core synthetic cells to hypertonic or hypotonic conditions by resuspending them in buffers with osmolarities equal to 200% or 50% that of the lumen, respectively (see **Synthetic cell swelling and shrinking** in the **Methods** for the buffer composition). Intriguingly, osmolarity-induced swelling and shrinking were seen to be more pronounced in gel-core synthetic cells compared with the liquid ones, with no apparent dependence on initial size observed (Figs. 2a-d, S5a-d, Movies S11-16).

The swelling and shrinking responses of hydrogels have been extensively characterised.^63^ When the external solute concentration is lowered relative to that of the gel-core lumen, water enters the lumen to dissipate the osmotic gradient. Hydration causes the PEG chains to expand, but because these are covalently linked in a continuous network, the polymers cannot diffuse apart, leading instead to macroscopic swelling of the gel. This response is absent in liquid-core synthetic cells, where unconstrained PEG molecules do not produce collective expansion. Conversely, increasing the external solute concentration drives water efflux and rapid hydrogel shrinking.

The membranes of gel-core synthetic cells could swell beyond their rupture limit, reported as a 3% increase in surface area,^64^ leading to membrane fissures of considerable size, as highlighted by the white arrows in Figures 2d,g and S5e. The 3D reconstruction of a swollen gel-core synthetic cell in Figure 2g further demonstrates membrane-fissure formation (yellow), exposing the gel core (red). These micron-scale defects are consistent with the cross-sectional synthetic-cell area increasing by approximately 20% (Figure S5c).

The large membrane fissures formed upon swelling of gel-core synthetic cells offer a valuable solution for controlling content exchange. Microscopy reveals leakage of encapsulated calcein from gel-core synthetic cells exposed to hypotonic conditions, with liquid-core synthetic cells remaining impermeable to the dye (Figs. 2e,g, Movies S17-19). Flow cytometry (FC) analysis confirms the complete loss of encapsulated calcein in populations of swollen synthetic cells with gel cores (Figs. 2h, S6), whereas those with liquid cores showed minimal calcein loss (Figs. 2f, S6).

We conducted a systematic study of the swelling-induced fissure formation as a function of osmolarity and (PEG-SH)_4_ concentration, testing gel-core synthetic cells with 2.5, 5.0, and 7.5 % w/v (PEG-SH)_4_ (Figs. 3a-b). Those with a lower mass fraction of (PEG-SH)_4_ were seen to swell and release encapsulated calcein at lower osmolarity mismatches (Fig. 3b). This trend is captured by a decreasing critical dilution factor (*DF_c_*) at which membrane fissure occurred, with increasing polymer density–an effect that could be leveraged to tune the permeabilization response in synthetic cells.

**Figure 3.**
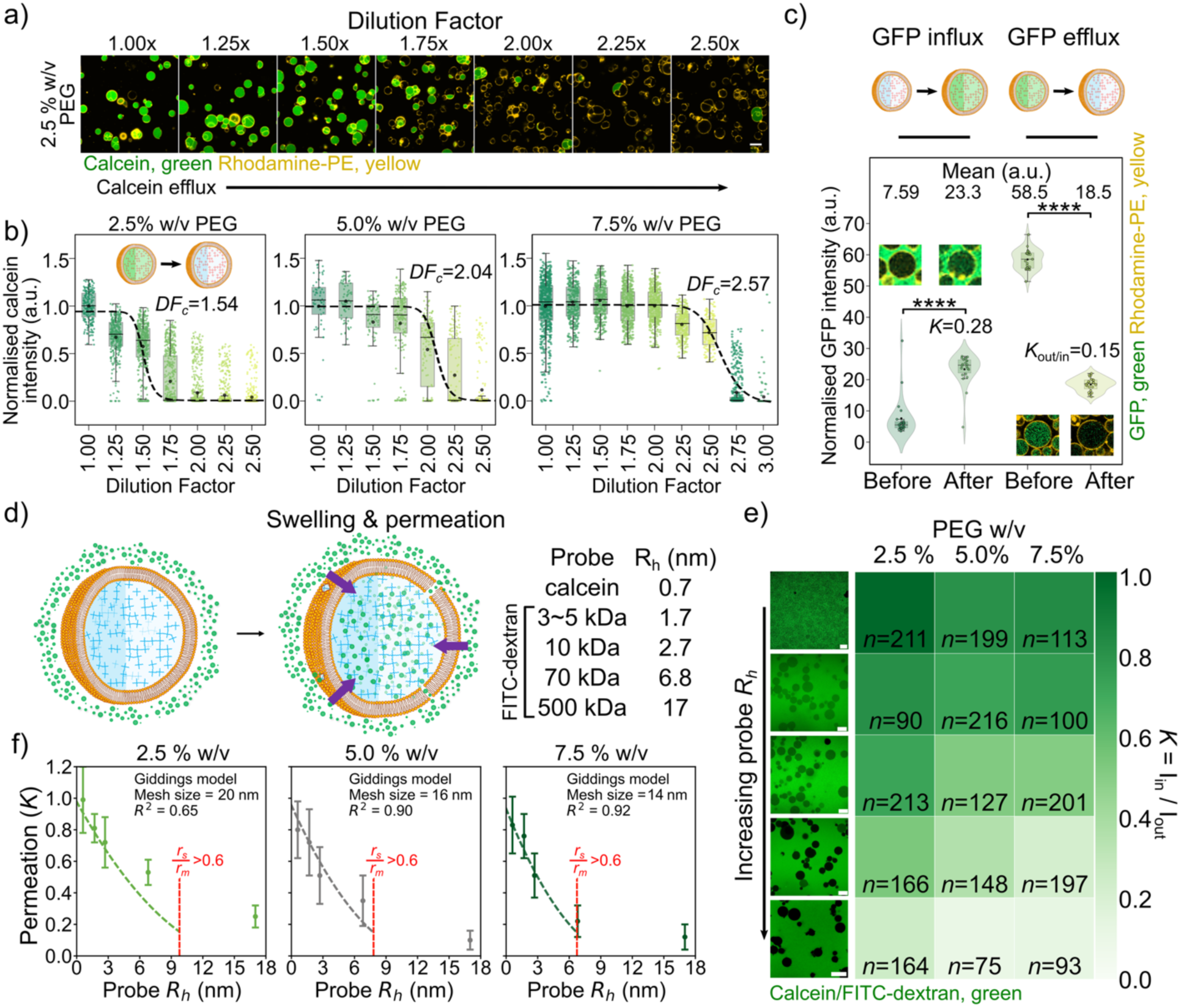
Membrane fissure formation in gel-core synthetic cells facilitates the influx and efflux of large macromolecules, with the extent of permeation dependent on the size of the molecule *R_h_* and gel mesh size (*r*_m_). (a) Confocal micrographs showing efflux of calcein (green) from gel-core synthetic cells encapsulating 2.5 % w/v (PEG-SH)_4_ under dilution into increasingly hypotonic conditions. Swelling of the gel core leads to membrane fissure formation and calcein release. The membrane is labelled with rhodamine-PE (yellow). (b) Box plots indicating calcein intensity in the lumen for gel-core synthetic cells initially encapsulating calcein, with (PEG-SH)_4_=2.5, 5.0 or 7.5% w/v. The calcein signal is normalized with 1 a.u. as the mean fluorescence intensity in initial isotonic conditions for a given (PEG-SH)_4_ concentration (*n* values provided in the SI). (c) Violin and box plots of the normalized intensity inside gel-core synthetic cells before and after swelling in hypotonic conditions (GFP-influx, *n*=22 and GFP-efflux, *n*=23). Inset confocal micrographs show exemplar synthetic cells in each condition, demonstrating GFP influx or efflux. *K* is defined generally as the fluorescent cargo partitioning into the synthetic cells (*K* = I_in_/I_out_); for efflux it is defined inversely as *K*_out/in_ = I_out_/I_in_. (d) Schematic of the permeation experiments of a green fluorophore into gel-core synthetic cells after swelling and membrane fissure formation. (e) Representative confocal micrographs at 5% w/v PEG showing the permeation of various fluorophores into swollen gel-core synthetic cells. The heat map indicates permeation (*K*), deduced from the relative fluorescence intensity of the synthetic cell interior (*I*_in_) versus the background intensity (*I*_out_) after 10 min incubation of the synthetic cells with various fluorescent molecules. (f) Line plots indicating the permeation of fluorescent probes into swollen gel-core synthetic cells as a function of probe size (*R_h_*) at varying (PEG-SH)_4_ concentrations. All data from (e) were plotted (mean ± 1 s.d.) and fit to the Giddings model using the theoretical mesh size (dashed line). The fits are shown up to *r_s_*/*r_m_* = 0.6 as the partitioning model is known to fail above this value, when the solute size is too large.^66^

Given the macroscopic size of the observed membrane fissures (Fig. 2g), we expected these to allow exchange of large macromolecular clients. Indeed, we observed the release of GFP (M_w_ 27 kDa) and OA647 from gel-core synthetic cells after hypotonic swelling (Figs. 3c, S7a, Movie S20). For both probes, ∼30% of the encapsulated proteins was retained (Fig. S8), likely due to cysteine-thiol linkages.^65^ We also confirmed that osmotic swelling causes uptake of GFP from the external solution in gel-core synthetic cells (Figs. 3c, S7b, Movie S21), but not in liquid-core synthetic cells (Fig. S9, Movie S22). These results confirm that swelling-induced membrane permeabilization can mediate the influx and efflux of proteins. Cysteine-mediated protein anchoring could be valuable to selectively hinder their release whilst exchanging other solutes; for instance, one could immobilize an enzyme in the gel and use swelling-induced permeabilization to exchange substrates and products. Where protein immobilization is undesirable, accessible cysteine residues may be blocked with maleimide derivatives.^64^

### Tunable permeability of gel-core synthetic cells

The possibility of tailoring the hydrogel mesh size, which we achieved by varying (PEG-SH)_4_ concentration, offers an additional means of controlling transport (Fig. 3d-e). We probed the partitioning of fluorescent probes with different sizes into swollen gel-core synthetic cells, using calcein (0.6 kDa)^67,68^ and various FITC-dextrans (3 to 500 kDa)^69^. The solute partitioning coefficient, *K,* was deduced from the relative fluorescence intensity of the synthetic cell interior (*I*_in_) versus the background unencapsulated material (*I*_out_), *i.e. K = I*_in_*/I*_out_. Higher *K* was seen with smaller molecular probes and reduced (PEG-SH)_4_ concentrations (Figs. 3e-f, Table S3).

The dashed lines in Figure 3f indicate predictions of *K* based on the Giddings model, which relates *K* to the polymer volume fraction (*φ*), solute radius (*r_s_*, here approximated as the hydrodynamic radius, *R_h_*) and gel mesh radius (*r_m_*).^70,71^ The theoretical estimates are in good agreement with the experimentally determined values of *K*._65,66_

Theoretical estimates of the hydrogel mesh radius (Supplementary Note 1) yielded 16, 12 and 11 nm for (PEG-SH)_4_ concentrations of 2.5, 5.0 and 7.5% w/v, respectively. These values are comparable to those obtained by nonlinear fitting of the partitioning coefficients using the Giddings model (31, 15 and 13 nm, respectively), indicating the validity of the models used. These observations suggest that solute permeability in the synthetic cells is regulated by hydrogel mesh size, including for macromolecules.

The possibility of controlling macromolecular transport through polymer concentration could prove beneficial for applications that involve the exchange of large payloads, including enzymes and transcription/translation machinery.^50,70^ We validated the expression of GFP in gel-core synthetic cells using the PUREfrex system (Fig. S10, Movies S23-24), indicating that the gel core does not disrupt relevant biomolecular reactions, consistent with prior reports.^51,72^

### Chemically tethering the gel core and inner leaflet enables membrane resealing

Having observed that interactions between the gelled lumen and the bilayer impact lipid diffusivity, we aimed to enhance the coupling between the gel core and lipid membrane via direct conjugation. This can be achieved by doping the membrane with maleimide-labelled phosphatidylethanolamine lipid (mal-PE), which can form covalent bonds with exposed thiols on the gel surface (Fig. 4a). The reactivity of mal-PE was first confirmed by encapsulating 0.1 % w/v AF647-PEG into synthetic cells prepared with or without mal-PE. We observed localization of AF647-PEG to the membrane when mal-PE was present (Fig. 4a), suggesting successful reaction between mal-PE and (PEG-SH)_4_, while the PEG-conjugated dye remained distributed throughout the lumen if mal-PE was absent (Fig. 4b). Furthermore, FRAP experiments demonstrated reduced mobility of the NBD-PE lipid in membranes containing mal-PE compared to those without it, supporting the inference that the mal-PE is anchored to (PEG-SH)_4_ (Fig. 1e).

**Figure 4.**
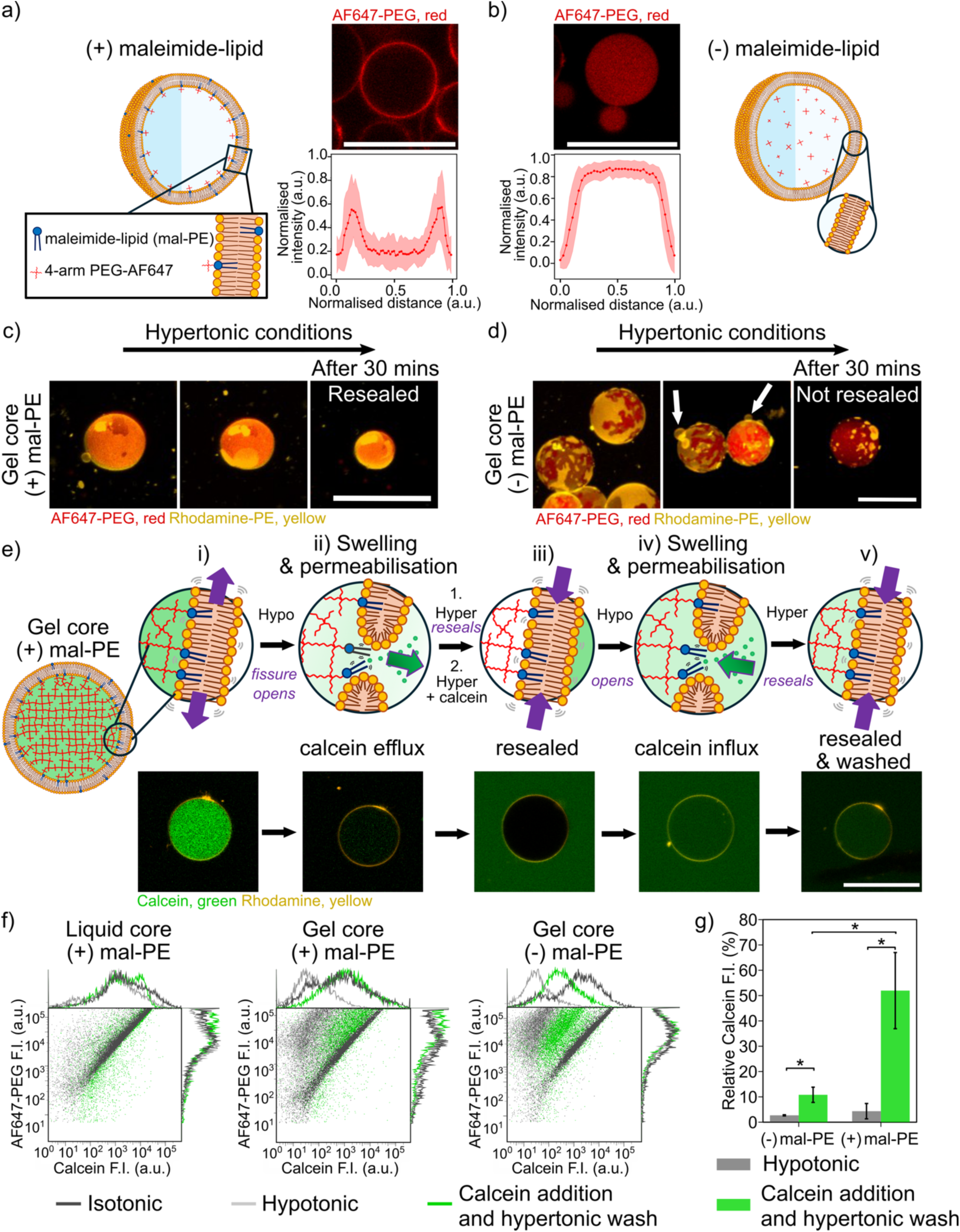
The gel can be directly conjugated to the membrane to facilitate membrane defect resealing. (a-b) Schematics indicating the location of AF647-PEG in liquid-core synthetic cells with (a) or without (b) mal-PE; confocal micrograph (middle top) and mean line plot profile (middle bottom, *n*=33) showing AF647-PEG fluorescence intensity across the synthetic cells in accordance with the schematics. Mal-PE was in vast excess to AF647-PEG (0.01%) in these experiments, resulting in the accumulation of all AF647 at the membrane in (a). Shaded regions in the line plots indicate ± 1 s.d. from the mean. The data are normalised by subtracting the background and scaling, with 1 being the maximum intensity in the synthetic cell population. (c-d) Confocal micrographs of swollen gel-core synthetic cells with (c) or without (d) mal-PE in the membrane. The gel cores are labelled with AF647 (red) and the membrane with rhodamine (yellow). Upon exposure to hypertonic conditions, the swollen gel-core synthetic cells shrink, with the membrane defects in (c) seen to reseal, whilst those in (d) do not. (e) Schematics (top) and associated confocal micrographs (bottom) indicating that hypotonic swelling of gel-core synthetic cells leads to content release (i, ii), with subsequent membrane resealing in hypertonic conditions in the presence of mal-PE (blue lipid) (iii). Addition of calcein externally shows no permeation until the synthetic cell is again swollen hypotonically (iv), with membrane resealing under a subsequent hypertonic wash (v) - overall facilitating content uptake and retention. (f) Two-dimensional FC dot plots of AF647-PEG *vs* calcein with histograms overlaid, for liquid- and gel-core (± mal-PE) synthetic cells initially encapsulating AF647-PEG and calcein (isotonic, dark grey), after exposure to hypotonic conditions (light grey), and after subsequent supplementation of calcein and a hypertonic wash (green) (*n*>10000 for all, see Figs. S11-12 for exact *n*). (g) Median calcein fluorescence intensity in gel-core synthetic cells before and after shrinking, taken from repeats of the FC experiments shown in (f) (*n*=3). All scale bars=20 µm. *p* < 0.05 (*).

We hypothesize that introducing a chemical linkage between the gel core and the inner membrane may facilitate resealing of the membrane fissures formed upon osmotic swelling, providing a route for reversible membrane permeabilization. We firstly subjected gel-core synthetic cells, either containing or lacking mal-PE, to hypotonic conditions to induce swelling and the formation of membrane fissures. These were then washed in hypertonic buffer to induce shrinking, and the membrane fissures were monitored (Fig. 4c). We observed fissure resealing in the presence of mal-PE, with the membrane (yellow) spreading back over the entire gel core (red). By contrast, in the absence of mal-PE, shrinking of the gel core led to the membrane crumpling and forming aggregates and blebs on the gel surface, leaving the fissures unrepaired (Fig. 4d).

We then tested whether the apparent resealing of the membrane translated into its diffusional barrier properties being restored. First, gel-core synthetic cells with mal-PE were made to swell and release encapsulated calcein (Fig. 4e (i,ii)). The released calcein was then washed away, and the synthetic cells shrunk using hypertonic conditions (Fig. 4e (iii)). Remarkably, calcein added to the external solution at this stage did not permeate the membrane, indicating resealing of the membrane fissures (Fig. 4e (iii)). We then again induced swelling of the synthetic cells, causing membrane fissure formation and calcein influx (Fig. 4e (iv). Finally, we resealed the membranes in hypotonic conditions and washed away unencapsulated calcein, ultimately observing the retention of the uptaken dye (Fig. 4e (v), Movie S25).

Monitoring calcein exchange, FC also confirmed the reversible formation and sealing of fissures in gel-core synthetic cells with mal-PE upon swelling/de-swelling (Fig. 4f, middle, Figs. S11). The same protocol applied to liquid-core synthetic cells (Fig. 4f, left, Fig. S12) resulted in minimal content exchange, while gel-core synthetic cells without mal-PE released but did not recapture calcein, indicating failure to reseal the fissures (Figs. 4f, right, Fig. S11). FC and LSCM analyses thus confirm that the presence of mal-PE is essential for the gel-core synthetic cells to exhibit cyclical content exchange.

Evidence that gel-bound lipids facilitate membrane resealing is intriguing. To the best of our knowledge, similar effects have not been previously reported, nor have hypotheses been put forward to clarify the underlying mechanism. We rationalize this observation as follows. When the gel swells, membrane fissures are introduced, leaving the acyl chains of the gel-bound lipids exposed in the areas not covered by the membrane (blue lipids in Figs. 4e, S13). When the gel shrinks back, there is a driving force for the membrane to re-wet these solvent-exposed areas due to the presence of the chemically bound lipids. This causes the membrane to spread back in its original configuration (Fig. S13, left). In contrast, in the absence of anchored lipids, there is no driving force for the membrane to re-wet the areas of the gel that become uncovered after swelling. The only driving force for membrane fissures to heal would therefore be that of minimizing the line energy of the fissures themselves, which is likely insufficient in this case to overcome energetic barriers associated with frictional forces (Fig. S13, right).

### External supply of molecular fuel powers intracellular biochemical reactions

Finally, we sought to verify the applicability of the cargo exchange method for controlling synthetic cell functionalities. In particular, we aimed to modulate internal biochemical reactions through the uptake of substrates, a critical capability that would enable synthetic cells to extract resources from the environment and extend their functional lifetime. Firstly, we designed gel-core synthetic cells (± mal-PE) encapsulating β-galactosidase (β-gal), which can hydrolyze non-fluorescent fluorescein di-β-D-galactopyranoside (FDG, 656 Da) into emissive fluorescein. β-gal is likely to be immobilized in the gel core due to cysteine-thiol bridging^73^, as observed for OA647 and GFP (Figs. S2, S3, S7-8). Control experiments confirm that such immobilization does not impact enzyme activity (Fig. S14). We proceeded to supply FDG in hypotonic conditions, aiming to induce membrane fissure formation and FDG uptake (Fig. 5a-ii), followed by the washing away of excess FDG in a hypertonic solution to reseal the fissures (Fig. 5a-iii), whilst monitoring the synthetic cell fluorescence over time (Fig. 5a-iv, v). After 60 min, gel-core synthetic cells (+)mal-PE showed a large increase in fluorescence, demonstrating FDG uptake, hydrolysis, and the retention of the fluorescein produced in their lumen (Fig. 5b, middle and Fig. 5c, Movie S26). Synthetic cells without mal-PE and liquid-core synthetic cells showed lower signals due to an inability to efficiently uptake the substrate and retain the product (Fig. 5,c, Movies S27-28). These results demonstrate that cyclic membrane permeabilization and resealing can be used to fuel internal biochemical reactions.

**Figure 5.**
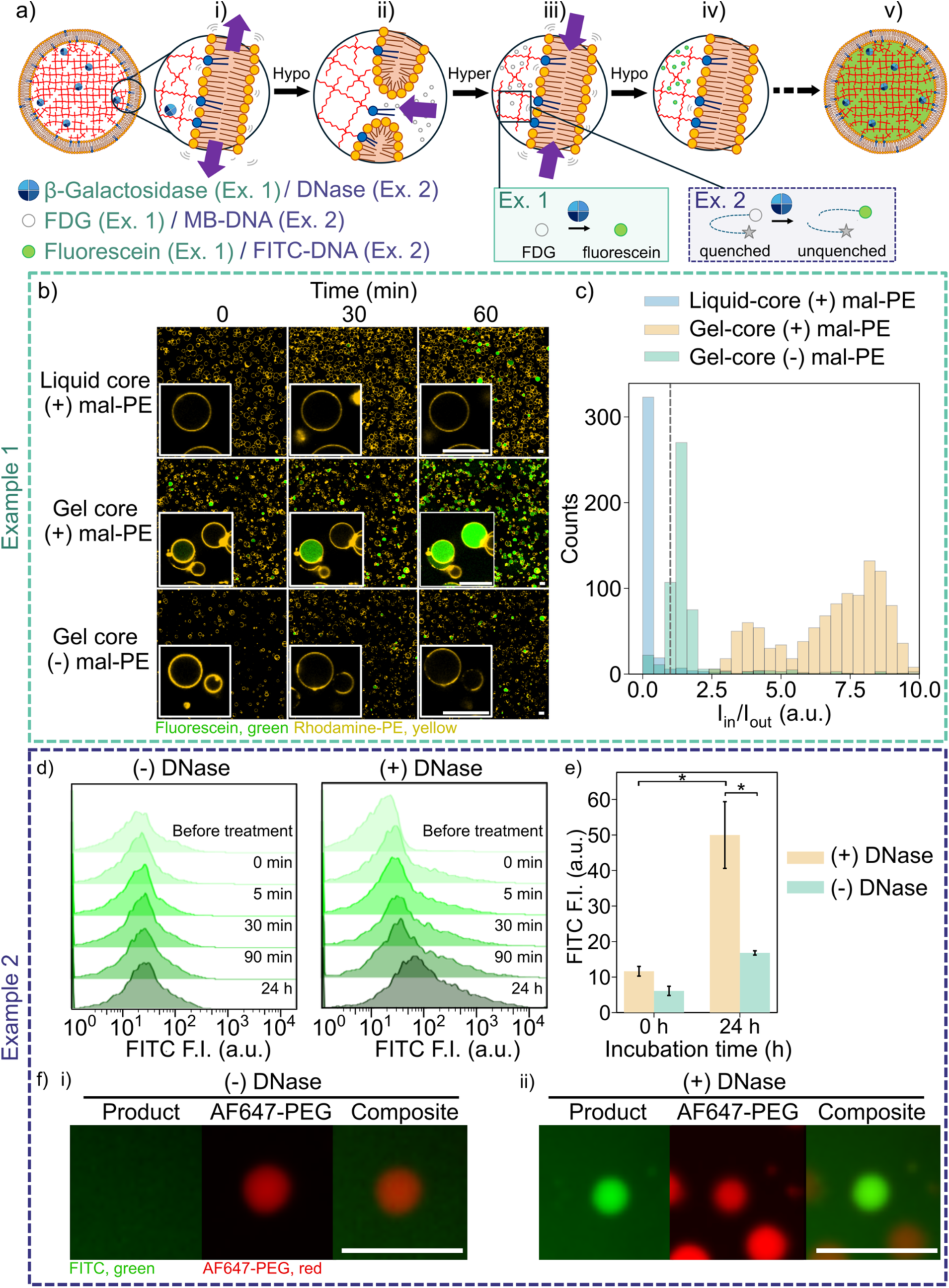
Membrane fissure formation and resealing facilitates substrate uptake and subsequent hydrolysis. (a) Schematics indicating that gel-core synthetic cells incorporating mal-PE (blue lipid) into the inner leaflet of the membrane are able to swell in hypotonic conditions with uptake of external FDG (Ex. 1) or molecular beacon-DNA (MB-DNA, Ex. 2); (i-ii), before membrane resealing and conversion of non-fluorescent starting material to the fluorescent product by either β-gal (Ex. 1) or DNase (Ex. 2) immobilized in the gel (iii-v). (b) Confocal micrographs showing the rhodamine and fluorescein channels overlaid, with representative synthetic cells in the inset (left, all scale bars=20 µm), and (c) associated histogram of the lumen fluorescence intensity (right) for liquid and gel (± mal-PE) synthetic cells exposed to the procedure shown in (a), i.e. incubation in hypotonic conditions, supplementation with FDG and subsequent incubation in hypertonic conditions. The histogram indicates the relative fluorescence intensity of the synthetic cell interior (*I_in_*) versus the background intensity (*I_out_*), with a line indicated at unity (*x*=1). (d) Histograms obtained via flow cytometry indicating the fluorescence intensity over time for gel-core synthetic cells (+mal-PE) without (left) and with (right) DNase co-encapsulation. In the presence of DNase, the fluorescence is seen to increase over time due to cleavage of the DNA molecular beacon, with (e) indicating that the median fluorescence intensity was higher than the control after 24 h. (f) Fluorescence micrographs of gel-core synthetic cells incorporating mal-PE after shrinking and resealing, after prior swelling in MB-containing hypotonic buffer. Scale bars=50 µm.

The exchange of proteins demonstrated that larger, macromolecular cargoes can be transported through the membrane fissures and hydrogel. We exploited this capability to fuel a second model biochemical reaction, whose substrate is a DNA molecular beacon (MB-DNA). The beacon consists of a hairpin structure featuring a fluorophore (FITC) and a quencher (BHQ-1) at the 5’ and 3’ends, respectively, leading to fluorescence quenching. Digestion by DNase can then restore fluorescent emission by distancing FITC from BHQ-1 (Fig. S15a-b). DNase was immobilized in the hydrogel through a maleimide-PEG4-NHS ester linker, as confirmed with a fluorescently-labelled version of the enzyme (Fig. S15c). We then supplied gel-core synthetic cells, loaded with immobilized (non-fluorescent) DNase, with external MB-DNA in a similar manner as the FDG/β-gal experiments, and analysed the fluorescence over time by FC and LSCM (Figs. 5d-f, S16). For enzyme-containing synthetic cells, FITC fluorescence increased around 5-fold during incubation, while those without DNase showed negligible signal. LSCM images were consistent with the FC data, confirming the successful supply of a (10.5 kDa) DNA fuel.

Our results thus confirm that the cyclic content-exchange mechanism we introduce can be used to modulate the biochemical activity of synthetic cells through the exchange of both small molecules and macromolecules, a critical capability that remains inaccessible with alternative approaches.

### Conclusions

We have demonstrated the design and construction of a previously unexploited class of composite synthetic cell, whereby responsive hydrogels are encapsulated within cell-sized GUVs. The hydrogel core affords the liposomes with an enhanced response to osmolarity changes, leading to shrinkage in hyperosmolar environments and swelling upon exposure to hypoosmolar solutions, thereby converting the hydrogel’s mechanical changes into membrane deformation. Expansion of the membrane upon gel swelling leads to fissure formation and the uptake of large macromolecules including proteins and DNA. Further engineering the lipid membrane’s inner leaflet to covalently bind the gel core enables resealing of the membrane fissures upon hyperosmolar shrinkage, thus establishing a synthetic pathway for cyclic and reversible membrane permeabilization. We deploy the pathway to build synthetic cells that can uptake a substrate, trigger an internal biochemical reaction, and retain the products following to efficient membrane resealing. Critically, the trafficking mechanism does not rely on biological protein channels or transporters, nor on synthetic nanopores, which are challenging to reconstitute and control in synthetic cellular devices. We therefore expect that our platform will substantially lower the technological barrier for engineering membrane trafficking in synthetic cells, potentially addressing key bottlenecks currently limiting the applicability of these devices as smart microreactors for healthcare and biomanufacturing. For instance, reliable solute exchange at specific timepoints could be used to periodically replenish ATP or other fuels to extend reaction lifetime, or to exchange functional macromolecules and dynamically alter the chemical profile of the synthetic cells for adaptive functions. Finally, one may speculate that protein-free membrane trafficking pathways that, like ours, are triggered by environmental oscillations, may have enabled content exchange in primordial cells, albeit with very different chemical details.

## Methods

### Chemicals

Horseradish peroxidase (HRP) (Cat. No. P8375; all following codes in brackets indicate the catalog number), hydrogen peroxide solution (H1009), fluorescein isothiocyanate– dextrans (FITC-dextrans) M_w_ 3000-5000 g mol^-1^ (FD4), average M_w_ 10,000 g mol^-1^ (FD10s), average M_w_ 70,000 g mol^-1^ (90718), average M_w_ 500,000 g mol^-1^ (FD500s), calcein (C0875), β-galactosidase from *Escherichia coli* (G5635), L-glutamic acid potassium salt monohydrate (KGlu) (G1149), HEPES (H3375), mineral oil (light) (M8410), 1-palmitoyl-2-{6-[(7-nitro-2-1,3-benzoxadiazol-4-yl)amino]hexanoyl}-sn-glycero-3-phosphoethanolamine) (NBD-PE) (810153C), 1,2-distearoyl-sn-glycero-3-phosphoethanolamine-N-[biotinyl(polyethylene glycol)-2000]) (DSPE-PEG(2000)-biotin) (880129P), 1-hexadecanoyl-2--(9Z-octadecenoyl)-sn-glycero-3-phosphocholine (POPC) (850457C) and 1,2-dioleoyl-sn-glycero-3-phosphoethanolamine-N-(lissamine rhodamine B sulfonyl) (ammonium salt) (rhodamine-PE) (810150C) were purchased from Merck. N-(3-Maleimide-1-oxopropyl)-L-α-phosphatidylethanolamine, dioleoyl (mal-PE) (COATSOME FE-8181MA3) and 4-arm thiolated PEG 20K ((PEG-SH)_4_) (SUNBRIGHT PTE-200SH) were purchased from NOF America. Alexa Fluor™ 647 C_2_ maleimide (10144342), Alexa Fluor™ 488 C_5_ maleimide (A10254), ovalbumin Alexa Fluor™ 647 conjugate (OA647) (O34784), glycyl-L-tyrosine hydrate (G01451G), dithiothreitol (DTT) (R0861), NeutrAvidin (neutravidin) (310000) and fluorescein Di-β-D-Galactopyranoside (FDG) (F1179) were purchased from Thermo Fisher Scientific. D-glucose anhydrous (0188-2.5KG) and D-sucrose (0335-2.5KG) were purchased from VWR Life Sciences. Magnesium acetate tetrahydrate (MgOAc) (139-15335) and DNase I (27-0516-01) were obtained from FUJIFILM Wako Chemicals.

GFP was purified from *E. coli* using pET-snap-GFP as described previously.^74^The ssDNA-based molecular beacon was synthesized by FASMAC (5’-FITC - TCTTGGACAAGTAAGGAGTCCATTGTGTCCAAGA -3’-BHQ).

### Synthetic cell preparation

Synthetic cells were formed using the GUV emulsion phase transfer method.^49,50^ Lipid mixtures were prepared from chloroform stocks (typically POPC: DSPE-PEG(2000)-biotin: rhodamine-PE at 98.5: 1: 0.5 mol %; FRAP experiments with POPC: NBD-PE at 99: 1 mol% (± 1 mol% mal-PE); maleimide binding experiments using POPC: DSPE-PEG(2000)-biotin at 99: 1 mol % (± 1 mol% mal-PE)) and 7.6 mg of the lipid mixture was deposited into the bottom of a glass tube. A gentle stream of nitrogen gas was used to evaporate the chloroform. Then, 0.5 ml of light mineral oil was added, heated at 70 °C and vortexed for 30 s repeatedly until the lipids were resuspended in the oil.

A water-in-oil (W/O) emulsion was prepared by vortexing for 30 s a mixture of 20 µL of inner buffer (19 mM HEPES-KOH pH 7.6, 241 mM KGlu, 17 mM MgOAc, 116 mM glucose, 270 mM sucrose, the indicated concentration of (PEG-SH)_4_, 5 mM glycy-l-tyrosine, ± 1 U µL^-1^ HRP) in 200 µL of the lipid-in-oil suspension. For the visualization of the synthetic cell core, (PEG-SH)_4_ was labelled with either AF488 or AF647 by mixing 1 mol% of AF488- or AF647-maleimide with 2.5 mM (PEG-SH)_4_ stock in Milli-Q, and this labelled (PEG-SH)_4_ was subsequently used for synthetic cell preparation. 10 µM calcein, 3.75 µM OA647, 10 µM GFP, 1 U µL^-1^ β-galactosidase or 0.1 mg mL^-1^ DNase I was added to the inner buffer as indicated in the text.

For the phase transfer, 200 µL of W/O emulsion was layered on top of 250 µl of outer buffer (20 mM HEPES-KOH, 250 mM KGlu, 18 mM MgOAc, and 400 mM glucose, pH 7.6) and centrifuged at 20 °C at 9000 *g* for 15 min. The oil layer was removed by pipetting and the GUV pellet (20 µL) collected, resuspended in 200 µL of outer buffer and centrifuged again at 20 °C at 6000 *g* for 5 min to wash away unencapsulated material. The washed pellet (20 µL) was collected and resuspended in 80 µL of outer buffer before further analysis. For GUVs containing mal-PE, 20 µL of a lipid-in-oil solution without mal-PE was layered on top of the outer buffer and settled for 30 min to facilitate the formation of the lipid monolayer at the water/oil interface. Then, the W/O emulsion with mal-PE was added on top of this layer, followed by centrifugation as described above.

To assess whether mal-PE in the membrane binds to encapsulated 4-arm PEG, liquid-core synthetic cells were prepared with 0.01 % w/v AF647-PEG ± 1 mol% mal-PE. These synthetic cells were added to a sample well placed on an imaging slide, sealed with a coverslip and single images were captured as shown in Figs. 4a,b.

### Synthetic cell swelling and shrinking

The initial external buffer and lumenal osmolarities are equal, at approximately 994 mOsmol/L, where the external buffer is identical to the outer buffer (20 mM HEPES-KOH, 250 mM KGlu, 18 mM MgOAc, and 400 mM glucose, pH 7.6) described above. Hypotonic conditions are induced by addition of an equivalent volume of water to the synthetic cell sample, to give an external buffer osmolarity of 0.5x the lumenal osmolarity (approx. 497 mOsmol/L), whilst hypertonic conditions are induced by adding an equivalent volume of outer buffer at 3x concentration to give an external osmolarity of 2x the lumenal osmolarity (approx. 1988 mOsmol/L). The external and lumenal osmolarities will re-equilibrate via synthetic cell swelling or pore formations. These steps can be combined sequentially to restore the initial osmolarity.

### Flowcytometric analysis

Flowcytometric (FC) analysis was carried out using a FACSAria III (BD Biosciences). For isotonic conditions, the liposome suspension was diluted 10-fold with outer buffer to a final volume of 200 µL and then subjected to FC analysis. At least 10,000 liposome particles were measured at a flow rate of ∼1000 events per second. Measurements under hypotonic conditions were carried out by 10-fold dilution with 0.5x outer buffer. Calcein was measured using a 488-nm blue laser and a 530/30 filter (434 V); rhodamine was measured using a 561-nm yellow laser and a 585/15 filter (300 V); AF647 was measured using a red 633 nm laser and a 660/20 filter (401 V). Thresholds were set at a value of 500 for both FSC and SSC, and 2500 for AF647. Data analysis was performed using FlowJo software (v10.9.0, BD Life Sciences).

### Confocal microscopy settings

Imaging of liposomes was carried out using either of the following confocal microscope configurations. The first comprised a Zeiss Axio Observer 7 fitted with an LSM 900 confocal scan head (LSM 900 with AiryScan 2, ZEISS) with a 40x/1.30 Plan Apo DIC (UV) VIS-IR oil immersion objective, and emission collected with MA-PMT detectors or an ESID for transmitted light. The second set-up comprised a Zeiss Axio Observer 7 fitted with an LSM 700 confocal scan head (LSM 700, ZEISS) with a 63x/1.40 Plan Apo DIC oil immersion objective and emission collected with GaAsP-PMT spectral detectors or a T-PMT for transmitted light. Fluorescent probes were detected between 490-nm and 555-nm (NBD, AF488; Ex 488-nm); 500-nm and 544-nm (calcein/fluorescein/FITC; Ex 494-nm); 560-nm and 630-nm (rhodamine; Ex 558-nm) and 655-nm and 700-nm (AF647; Ex 653-nm) using the relevant detectors. Calcein/fluorescein/FITC/AF488/GFP and AF647 signals were simultaneously acquired in channel mode with unidirectional scanning and switching track every frame, using the 488-nm and 647-nm diode laser lines on the same track as transmitted light (if used), whilst rhodamine signals were acquired on a second channel using the 561-nm diode laser line, with all lasers set to 2% using the Zen software unless stated otherwise. The images were focused on a single plane near the bottom of the glass slide where vesicles had settled and imaged using Leica Microsystems Immersion Oil and a 1 Airy unit pinhole unless otherwise specified. Zen (blue edition) acquisition software version 3.5 (ZEISS) was used to acquire the data. Images were analysed and ROI measurements extracted using Fiji software.^75^ The extracted image data was subsequently analysed using Python (see “**Data Processing**”).

Unless noted, synthetic cells were prepared with 1% biotinylated lipid (DSPE-PEG2000-biotin) and imaged on neutravidin and BSA-coated glass imaging slides with a thickness of 0.13-0.17 mm to enable immobilization via biotin-streptavidin interactions. For the coating, 1% BSA and 100 nM neutravidin, respectively, were dissolved in outer buffer. The slides were first incubated with BSA for 1 h and subsequently neutravidin solution for another 1 h, followed by rinsing with Milli-Q and drying with nitrogen gas. Coated slides were kept at 4°C until use. Vesicles were imaged in Φ1x1 mm silicon or PDMS wells placed on the coated glass slides and sealed with a coverslip. When supplying osmolytes or reagents during imaging, the synthetic cell solution was first loaded on the glass slide, and the additives were subsequently supplied by gentle pipetting onto the top of the imaging chamber.

### Confirming the gelation of the hydrogel in the synthetic cell lumen

Time-lapse imaging was performed before and after bleaching with a 1.63 s interval (Figs. 1b-c, S2, S4, S6) or 1 s interval (Fig. S3). 3 images were collected before photobleaching. Photobleaching was carried out at 50% of the 488-nm or 640-nm lasers in the ROI (bleach duration = 500 ms; 10 iterations).

To image the gel-to-liquid transition after adding DTT to gel-core synthetic cells (Fig. S4), the gel core was first bleached, before the addition of DTT (5 mM, in isotonic conditions). Time-lapse imaging was then performed with a 10 ms interval (imaging condition identical to that for photobleaching except pixel dwell time: 1.50 µs). To image synthetic cell lysis and content release (Fig. S1), Triton-X-100 was added (2 % v/v, in isotonic buffer) before covering the sample with a coverslip. Time-lapse imaging was performed with a 10 s interval (imaging condition identical to that for photobleaching). FC measurements after detergent lysis (Fig. 2e-h) were performed by first supplementing the synthetic cells with Triton-X-100 (2 % v/v, in isotonic buffer) and incubating for 10 min, before dilution and FC measurement.

### Synthetic cell permeability at varying osmolarities, PEG concentrations and probe sizes

Flory-Rehner theory describes gel swelling as a balance between favourable interactions between the polymer chains and solvent, and the elastic tension generated by stretching of the polymer chains.^76^ By diluting samples of gel-core synthetic cells into Milli-Q, the osmolarity of the buffer was sequentially reduced, and the calcein leakage via induced membrane fissures was measured for synthetic cells with different concentrations of (PEG-SH)4. In brief, gel-core synthetic cells were prepared with 2.5, 5.0 or 7.5 % w/v AF647-labelled (PEG-SH)_4_, 10 μM calcein and 0.5 mol % rhodamine-PE, and incubated in hypotonic conditions by adding Milli-Q to the synthetic cell sample at the ratios of synthetic cell sample to Milli-Q shown in Table S5. The fluorescence intensity of the gel-core synthetic cells reduced when calcein leaked out. By fitting a sigmoidal curve to the calcein fluorescence intensity (Fig. 3b), the dilution factor (*DF*_C_) was estimated.

To reveal the relationship between the synthetic cell permeability and the probe size, synthetic cells were placed in hypotonic conditions by adding an equivalent volume of Milli-Q to a given volume of synthetic cell sample, before addition of 5 µM of the fluorescent probe (calcein or 3-5, 10, 70 or 500 kDa FITC-dextran) and incubation for 10 min at room temperature. Images as shown in Figure 3e were then captured post-incubation.

The relationship between the solute partitioning coefficient, *K,* and the hydrodynamic radius, *R_h_,* of the various fluorescent probes was plotted using the Giddings model (Eq. 1), which relates *K* with the polymer volume fraction (*φ*), solute radius (*r_s_*, here approximated as *R_h_*) and mesh radius (*r_m_*).

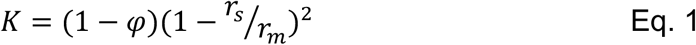

This partitioning model is known to fail when the solute size is too large, when *r_s_*/*r_m_* > 0.6.^66^ The encapsulated PEG concentration and PEG dry density (*ρ_d_* = 1.18 g mL^-1^) was used to calculate *φ*:

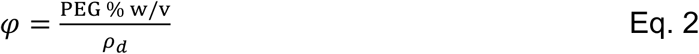

To fit the Giddings model, *r_m_* was calculated based on the theoretically estimated mesh sizes (see **Supplementary Note 1**), and *r_s_* was set as the probe hydrodynamic radius (*rs* = *R_h_)*. The *r_m_* was also obtained by fitting Eq.1 to the data shown in Fig. 3f.

### Calcein release and subsequent encapsulation

For measurements of calcein efflux and influx (Fig. 4e,f,g), gel- or liquid-core synthetic cells (± 1 mol% mal-PE) encapsulating 10 µM calcein were prepared. Biotinylated lipid was not included in these samples to eliminate the possibility that interactions between the glass slide and the lipid membrane prevent efficient membrane resealing. Calcein release and subsequent uptake was measured by subjecting synthetic cells to the following sequence: (i) 10 min incubation in solution diluted with equivalent volume of Milli-Q (osmolarity = 0.5x initial osmolarity), causing calcein release; (ii) addition of 3x outer buffer, causing gel-core synthetic cell shrinking and membrane resealing (osmolarity = 1x initial osmolarity) (iii) 10 min incubation with 10 µM calcein in outer buffer; (iv) addition of 10 µM calcein in Milli-Q, leading to calcein influx (osmolarity = 0.5x initial osmolarity); (v) 10 min incubation in <u>3</u>x outer buffer containing 10 µM calcein (osmolarity = 1x initial osmolarity); (vi) addition of outer buffer (osmolarity = 1x initial osmolarity) to give a final osmolarity equivalent to the initial outer buffer. Time-lapse imaging of the same sequence (Fig. 4c, Movie S22) was performed for 51 min with a 10 s interval.

### Assessment of membrane fluidity and molecule diffusion in the gel network using FRAP

ROIs of size 1.44 µm x 1.52 µm were used for membrane bleach experiments of NBD-PE. Circular ROIs of radius 2.12 µm^2^ were used for bleach experiments of FITC-dextran in the gel core. The ROI was bleached for 50 (NBD-PE) or 100 (FITC-dextran) iterations at 100% laser power using the 488 nm laser. Emission was collected for the ROIs at the appropriate wavelength for each fluorescence compound. Time-lapse imaging (Fig. 1d) was performed for 60 frames with an interval of 614.4 ms (NBD-PE) or 100 frames with an interval of 34.9 ms (FITC-dextran). 20 (NBD-PE) or 10 (FITC-dextran) images were collected pre-bleaching and 40 (NBD-PE) or 90 (FITC-dextran) post-bleaching at 1% laser power. Synthetic cells with large membrane fluctuations, fluorescent aggregates in the ROI, or that moved during acquisition were excluded from analysis. In order to record the fast bleaching and recovery dynamics of FITC-dextran in the gel core, only the bleaching ROI was acquired, as this speeds up the frame acquisition time considerably.

### Calculation of *t_1/2_* and diffusion coefficients from the FRAP data

FRAP data fitting was performed using Python after extraction of fluorescence intensity data using Fiji. We obtained *I*(*t*)_bleach_, the fluorescence intensity in the region with fluorescence that undergoes targeted bleaching, *I*(*t*)_non-bleach_, the fluorescence intensity in a different region with fluorescence in the same synthetic cell that does not undergo targeted bleaching, and *I*(*t*)_bk_, the fluorescence intensity in the background ROI that does not have fluorescence and does not undergo targeted bleaching. The images for all were acquired at 1% laser intensity. Background subtraction was performed using:

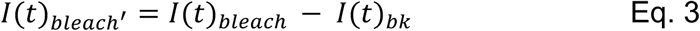

Photobleaching normalization was subsequently carried out by first fitting an exponential decay to the non-bleached ROI data:^77^

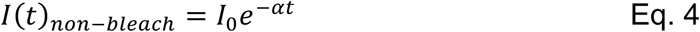

Where *I*(*t*) is the fluorescence at a given time, *t*, *I*_0_ is the initial fluorescence, and *α* is the decay constant. The bleach ROI *I*(*t*)*_bleac_*_ℎ_′ was then scaled first by *e*^−*αt*^, scaled, and second with the average of the first 20 frames (pre-bleach) set to 1 and the intensity immediately post-bleach to 0 to obtain *I*(*t*)*_bleac_*_ℎ_′__*norm*_. This normalised data was then fit to the following equation:

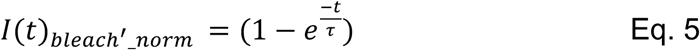

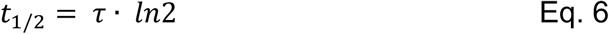

Where τ is the relaxation time and *t*_1/2_is the half maximal recovery time. Finally, the apparent diffusion coefficient *D_app_* was calculated as follows:^78,79^

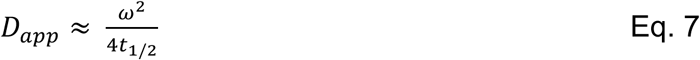

Where *ω* is the radius of the observation ROI (= 1.44 µm). Although the FRAP measurements were bleached and acquired within a square ROI for the NBD-PE data, the diameter of the bleaching spot in this case is similar to the z-resolution of the confocal (∼1 µm), and therefore we approximated the apparent diffusion coefficient using the formula in Eq. 5 for a circular bleaching spot. The calculated *D_app_* values and previously reported values are described as mean ± mean squared error (MSE).

### Enzymatic reaction in the synthetic cell lumen using substrates supplied from outside

Synthetic cells were prepared with β-gal (1U µL ^-1^) and added to an imaging well. Synthetic cells were treated with 0.75 mM FDG in hypotonic conditions and incubated for 10 min before changing to hypertonic conditions. Time-lapse imaging (Fig. 5b) was performed for 1 h with a 3 min interval.

DNase I does not have Cys residues and thus was labelled with TAMRA as follows. 1.5 mM mal-PEG_4_-NHS was premixed with 1.8 mM TAMRA-thiol in 16.5 µL DMSO and incubated for 2 h at room temperature. Together with the reaction mix, 1.25 mg of DNase was added to 200 µL of 10 mM HEPES, 100 mM NaCl buffer (pH7.4) at room temperature for 2 h to produce TAMRA-labeled DNase. The reaction mixture was dialyzed against 10 mM HEPES and 100 mM NaCl for 2 days at 4°C with continuous stirring and concentrated with an Amicon ultracentrifuge filter (MWCO 10kDa). To prepare gel-core synthetic cells encapsulating 0.1 mg/mL DNase or 0.05 mg/mL TAMRA-labeled DNase, 10 µM mal-PEG4-NHS was premixed with DNase in the buffer containing 5 mM HEPES and 50 mM NaCl and incubated for 2 h at room temperature. Subsequently, 5% (PEG-SH)_4_ was added to the mixture and incubated for another 2 h at room temperature, followed by the addition of other inner buffer components and used for gel-core synthetic cell formation as indicated in the synthetic cell preparation section. To supply MB DNA from outside the synthetic cells, experiments were done via the same process as with FDG, except that FDG was replaced with 15 µM MB DNA and incubated for 2 min before resealing. The time-course of fluorescein intensity was monitored by BD FACS Aria III, and the microscopic images were taken using Leica Epifluorescence microscope DMI6000B with 120 W mercury lamp.

### Data Processing and Statistical Analysis

The data was processed using Fiji^75^ and Python. The Student’s *t*-test (unpaired, two-tailed; using scipy.stats.ttest_ind) was used for comparative analysis. *p* < 0.05 was considered statistically significant. Explicit *p* values for the relevant Figures are shown in Table S6. In all violin-plots and box-plots (Figs. 1e-f; 3b-c; S12, S13) the centre line depicts the median and the boxes span the interquartile range (25–75%); when present, the black dot indicates the mean.

## Data Availability

Raw data supplied.

## Acknowledgements

We thank Takayoshi Watanabe for preparing the GFP used in this study. This work was supported by a JSPS Postdoctoral Fellowship for Research in Japan (short-term) (A.C.) and Leverhulme Trust (Doctoral Scholarship 2020) (A.C.), Human Frontier Science Program Grant Number RGP003/2023 (T.M.), JSPS KAKENHI Grant Numbers 22K21344 (T.M.), 21H05228 (T.M.), JST ASPIRE Grant Number JPMJAP24B4 (T.M.), BBSRC Grant Number BB/X012565/1 (L.D.M., Y.E.) and International Science Partnerships Fund (ISPF) Project ref. 250 (L.D.M, Y.E., T.M.).

## Author Contributions

A.C. and T.M. conceived and designed the experiments. A.C. and W.Z. performed the experiments. A.C, wrote the original draft with review and editing from W.Z., L.D.M, Y.E. and T.M. Funding was acquired by A.C., L.D.M, Y.E. and T.M. Research was supervised and directed by L.D.M, Y.E. and T.M.

## Notes

### Competing Interest Statement

The authors have declared no competing interest.

### Summary of Updates

Abstract and Introduction modified for clarity. Fig. 3 and 5 have been revised. The supplemental file has been updated.

## References

1. Vu, T. Q., Sant’Anna, L. E. & Kamat, N. P. Tuning Targeted Liposome Avidity to Cells via Lipid Phase Separation. Biomacromolecules 24, 1574–1584 (2023).

2. Sato, Y. & Takinoue, M. Capsule-like DNA Hydrogels with Patterns Formed by Lateral Phase Separation of DNA Nanostructures. JACS Au 2, 159–168 (2022).

3. Stewart, J. M. et al. Modular RNA motifs for orthogonal phase separated compartments. Nat. Commun. 15, 6244 (2024).

4. Noba, K. et al. Simple Method for the Creation of a Bacteria-Sized Unilamellar Liposome with Different Proteins Localized to the Respective Sides of the Membrane. ACS Synth. Biol. 12, 1437–1446 (2023).

5. Wang, M., Liu, Z. & Zhan, W. Janus Liposomes: Gel-Assisted Formation and Bioaffinity-Directed Clustering. Langmuir 34, 7509–7518 (2018).

6. Izri, Z., Garenne, D., Noireaux, V. & Maeda, Y. T. Gene Expression in on-Chip Membrane-Bound Artificial Cells. ACS Synth. Biol. 8, 1705–1712 (2019).

7. Aufinger, L. & Simmel, F. C. Artificial Gel-Based Organelles for Spatial Organization of Cell-Free Gene Expression Reactions. Angew. Chem. Int. Ed. 57, 17245–17248 (2018).

8. Adamala, K. P., Martin-Alarcon, D. A., Guthrie-Honea, K. R. & Boyden, E. S. Engineering genetic circuit interactions within and between synthetic minimal cells. Nat. Chem. 9, 431 (2017).

9. Niederholtmeyer, H., Chaggan, C. & Devaraj, N. K. Communication and quorum sensing in non-living mimics of eukaryotic cells. Nat. Commun. 9, 5027 (2018).

10. Ishii, Y., Fukunaga, K., Cooney, A., Yokobayashi, Y. & Matsuura, T. Switchable and orthogonal gene expression control inside artificial cells by synthetic riboswitches. Chemical Communications 60, 5972–5975 (2024).

11. Berhanu, S., Ueda, T. & Kuruma, Y. Artificial photosynthetic cell producing energy for protein synthesis. Nat. Commun. 10, 1–10 (2019).

12. Park, D. et al. Motility analysis of bacteria-based microrobot (bacteriobot) using chemical gradient microchamber. Biotechnol. Bioeng. 111, 134–143 (2014).

13. Kumar, B. V. V. S. P., Patil, A. J. & Mann, S. Enzyme-powered motility in buoyant organoclay/DNA protocells. Nat. Chem. 10, 1154–1163 (2018).

14. Siton-Mendelson, O. & Bernheim-Groswasser, A. Toward the reconstitution of synthetic cell motility. Cell Adhesion and Migration vol. 10 461–474 Preprint at 10.1080/19336918.2016.1170260 (2016).

15. Gispert, I. et al. Stimuli-responsive vesicles as distributed artificial organelles for bacterial activation. Proceedings of the National Academy of Sciences 119, (2022).

16. Smith, J. M., Chowdhry, R. & Booth, M. J. Controlling Synthetic Cell-Cell Communication. Front. Mol. Biosci. 8, (2022).

17. Mukwaya, V., Mann, S. & Dou, H. Chemical communication at the synthetic cell/living cell interface. Commun. Chem. 4, 161 (2021).

18. Niederholtmeyer, H., Chaggan, C. & Devaraj, N. K. Communication and quorum sensing in non-living mimics of eukaryotic cells. Nat. Commun. 9, 1–8 (2018).

19. Fujii, S. et al. Liposome display for in vitro selection and evolution of membrane proteins. Nat. Protoc. 9, 1578–1591 (2014).

20. Hartmann, D., Chowdhry, R., Smith, J. M. & Booth, M. J. Orthogonal Light-Activated DNA for Patterned Biocomputing within Synthetic Cells. J. Am. Chem. Soc. 145, 9471–9480 (2023).

21. Sampson, K., Sorenson, C. & Adamala, K. P. Preparing for the future of precision medicine: synthetic cell drug regulation. Synth. Biol. 9, (2024).

22. Lopez, A. & Fiore, M. Investigating Prebiotic Protocells for a Comprehensive Understanding of the Origins of Life: A Prebiotic Systems Chemistry Perspective. 9, 49 (2019).

23. van Dongen, S. F. M. et al. Cellular Integration of an Enzyme-Loaded Polymersome Nanoreactor. Angewandte Chemie 122, 7371–7374 (2010).

24. Discher, B. M. et al. Polymersomes: Tough vesicles made from diblock copolymers. Science (1979). 284, 1143–1146 (1999).

25. Akashi, K. I., Miyata, H., Itoh, H. & Kinosita, K. Preparation of giant liposomes in physiological conditions and their characterization under an optical microscope. Biophys. J. 71, 3242–3250 (1996).

26. Fujii, S. et al. Purification-Free MicroRNA Detection by Using Magnetically Immobilized Nanopores on Liposome Membrane. Anal. Chem. 90, 10217–10222 (2018).

27. Ugrinic, M. et al. Microfluidic formation of proteinosomes. Chemical Communications 54, 287–290 (2018).

28. Huang, X., Patil, A. J., Li, M. & Mann, S. Design and Construction of Higher-Order Structure and Function in Proteinosome-Based Protocells. J. Am. Chem. Soc. 136, 9225–9234 (2014).

29. Wen, P. et al. Construction of Eukaryotic Cell Biomimetics: Hierarchical Polymersomes-in-Proteinosome Multicompartment with Enzymatic Reactions Modulated Protein Transportation. Small 17, (2021).

30. Garenne, D. et al. Sequestration of Proteins by Fatty Acid Coacervates for Their Encapsulation within Vesicles. Angewandte Chemie 128, 13673–13677 (2016).

31. Poudyal, R. R. et al. Template-directed RNA polymerization and enhanced ribozyme catalysis inside membraneless compartments formed by coacervates. Nat. Commun. 10, 490 (2019).

32. Dai, Y. et al. Programmable synthetic biomolecular condensates for cellular control. Nat. Chem. Biol. 19, 518–528 (2023).

33. Laos, R. & Benner, S. Fluorinated oil-surfactant mixtures with the density of water: Artificial cells for synthetic biology. PLoS One 17, e0252361 (2022).

34. Torre, P., Keating, C. D. & Mansy, S. S. Multiphase Water-in-Oil Emulsion Droplets for Cell-Free Transcription–Translation. Langmuir 30, 5695–5699 (2014).

35. Kahn, J. S. et al. DNA Microgels as a Platform for Cell-Free Protein Expression and Display. Biomacromolecules 17, 2019–2026 (2016).

36. Wanselius, M., Rodler, A., Searle, S. S., Abrahmsén-Alami, S. & Hansson, P. Responsive Hyaluronic Acid–Ethylacrylamide Microgels Fabricated Using Microfluidics Technique. Gels 8, 588 (2022).

37. Allen, M. E., Hindley, J. W., Law, R. V., Ces, O. & Elani, Y. Microfluidic Production of Spatially Structured Biomimetic Microgels as Compartmentalized Artificial Cells. Small Science 5, 2400320 (2025).

38. Jørgensen, I. L., Kemmer, G. C. & Pomorski, T. G. Membrane protein reconstitution into giant unilamellar vesicles: a review on current techniques. European Biophysics Journal 2016 46:2 46, 103–119 (2016).

39. Fan, S. et al. Morphology remodelling and membrane channel formation in synthetic cells via reconfigurable DNA nanorafts. Nat. Mater. 24, 278–286 (2025).

40. Noireaux, V. & Libchaber, A. A vesicle bioreactor as a step toward an artificial cell assembly. Proc. Natl. Acad. Sci. U. S. A. 101, 17669–17674 (2004).

41. Hilburger, C. E., Jacobs, M. L., Lewis, K. R., Peruzzi, J. A. & Kamat, N. P. Controlling Secretion in Artificial Cells with a Membrane and Gate. ACS Synth. Biol. 8, 1224–1230 (2019).

42. Wang, X. et al. ON–OFF nanopores for optical control of transmembrane ionic communication. Nature Nanotechnology 2025 20:3 20, 432–440 (2025).

43. Fan, S. et al. A synthetic cell microreactor with two types of interacting dynamic DNA-based pores. Nature Chemistry 2026 18:7 18, 1325–1333 (2026).

44. Dey, S. et al. A reversibly gated protein-transporting membrane channel made of DNA. Nature Communications 2022 13:1 13, 2271-(2022).

45. Li, D. Y., Zhou, Z. H., Yu, Y. L. & Deng, N. N. Microfluidic construction of cytoskeleton-like hydrogel matrix for stabilizing artificial cells. Chem. Eng. Sci. 264, 118186 (2022).

46. Mou, Y., Zhang, P., Lai, W. F. & Zhang, D. Design and applications of liposome-in-gel as carriers for cancer therapy. Drug Deliv. 29, 3245–3255 (2022).

47. Llopis-Lorente, A. et al. Artificial cells with viscoadaptive behavior based on hydrogel-loaded giant unilamellar vesicles. Chem. Sci. 15, 629–638 (2024).

48. Shibayama, M. Physics of polymer gels: Toyoichi Tanaka and after. Soft Matter 21, 1995– 2009 (2025).

49. Pautot, S., Frisken, B. J. & Weitz, D. A. Production of Unilamellar Vesicles Using an Inverted Emulsion. Langmuir 19, 2870–2879 (2003).

50. Nishimura, K. et al. Population Analysis of Structural Properties of Giant Liposomes by Flow Cytometry. Langmuir 25, 10439–10443 (2009).

51. Kamiya, N., Ohama, Y., Minamihata, K., Wakabayashi, R. & Goto, M. Liquid Marbles as an Easy-to-Handle Compartment for Cell-Free Synthesis and In Situ Immobilization of Recombinant Proteins. Biotechnol. J. 13, 1–5 (2018).

52. Grzybek, M., Kubiak, J., Łach, A., Przybyło, M. & Sikorski, A. F. A Raft-Associated Species of Phosphatidylethanolamine Interacts with Cholesterol Comparably to Sphingomyelin. A Langmuir-Blodgett Monolayer Study. PLoS One 4, e5053 (2009).

53. Schaich, M., Sobota, D., Sleath, H., Cama, J. & Keyser, U. F. Characterization of lipid composition and diffusivity in OLA generated vesicles. Biochimica et Biophysica Acta (BBA) - Biomembranes 1862, 183359 (2020).

54. Guo, L. et al. Molecular Diffusion Measurement in Lipid Bilayers over Wide Concentration Ranges: A Comparative Study. ChemPhysChem 9, 721–728 (2008).

55. Przybylo, M. et al. Lipid Diffusion in Giant Unilamellar Vesicles Is More than 2 Times Faster than in Supported Phospholipid Bilayers under Identical Conditions. Langmuir 22, 9096– 9099 (2006).

56. Renner, L. et al. Supported Lipid Bilayers on Spacious and pH-Responsive Polymer Cushions with Varied Hydrophilicity. J. Phys. Chem. B 112, 6373–6378 (2008).

57. Axpe, E. et al. A Multiscale Model for Solute Diffusion in Hydrogels. Macromolecules 52, 6889–6897 (2019).

58. Richbourg, N. R. & Peppas, N. A. Solute diffusion and partitioning in multi-arm poly(ethylene glycol) hydrogels. J. Mater. Chem. B 11, 377–388 (2023).

59. Stein, P. E., Leslie, A. G. W., Finch, J. T. & Carrell, R. W. Crystal structure of uncleaved ovalbumin at 1·95 Å resolution. J. Mol. Biol. 221, 941–959 (1991).

60. Campanella, B., Onor, M., D’Ulivo, A., Giannarelli, S. & Bramanti, E. Impact of Protein Concentration on the Determination of Thiolic Groups of Ovalbumin: A Size Exclusion Chromatography–Chemical Vapor Generation–Atomic Fluorescence Spectrometry Study via Mercury Labeling. Anal. Chem. 86, 2251–2256 (2014).

61. Wakabayashi, R., Ramadhan, W., Moriyama, K., Goto, M. & Kamiya, N. Poly(ethylene glycol)-based biofunctional hydrogels mediated by peroxidase-catalyzed cross-linking reactions. Polym. J. 52, 899–911 (2020).

62. Yu, W. et al. Synthesis of functional protein in liposome. J. Biosci. Bioeng. 92, 590–593 (2001).

63. Feng, W. & Wang, Z. Tailoring the Swelling-Shrinkable Behavior of Hydrogels for Biomedical Applications. Advanced Science 10, 2303326 (2023).

64. Koslov, M. M. & Markin, V. S. A theory of osmotic lysis of lipid vesicles. J. Theor. Biol. 109, 17–39 (1984).

65. Yang, F., Moss, L. G. & Phillips, G. N. The Molecular Structure of Green Fluorescent Protein. Nat. Biotechnol. 14, 1246–1251 (1996).

66. Cu, Y. & Saltzman, W. M. Mathematical modeling of molecular diffusion through mucus. Adv. Drug Deliv. Rev. 61, 101 (2008).

67. Tamba, Y., Ariyama, H., Levadny, V. & Yamazaki, M. Kinetic pathway of antimicrobial peptide magainin 2-induced pore formation in lipid membranes. Journal of Physical Chemistry B 114, 12018–12026 (2010).

68. Chenyakin, Y. et al. Diffusion coefficients of organic molecules in sucrose-water solutions and comparison with Stokes-Einstein predictions. Atmos. Chem. Phys. 17, 2423–2435 (2017).

69. Hanselmann, R., Burchard, W., Lemmes, R. & Schwengers, D. Characterization of DEAE-dextran by means of light scattering and combined size-exclusion chromatography/low-angle laser light scattering/viscometry. Macromol. Chem. Phys. 196, 2259–2275 (1995).

70. Giddings, J. C., Kucera, E., Russell, C. P. & Myers, M. N. Statistical theory for the equilibrium distribution of rigid molecules in inert porous networks. Exclusion chromatography. Journal of Physical Chemistry 72, 4397–4408 (1968).

71. Pluen, A., Netti, P. A., Jain, R. K. & Berk, D. A. Diffusion of Macromolecules in Agarose Gels: Comparison of Linear and Globular Configurations. Biophys. J. 77, 542–552 (1999).

72. Koga, T. et al. A High-Throughput Cell-Free Enzyme Screening System Using Redox-Responsive Hydrogel Beads as Artificial Compartments. ACS Synth. Biol. 14, 995–1001 (2025).

73. Jornvall, H., Fowler, A. V. & Zabin, I. Probe of β-galactosidase structure with iodoacetate. Differential reactivity of thiol groups in wild-type and mutant forms of β-galactosidase. Biochemistry 17, 5160–5164 (1978).

74. Uyeda, A., Watanabe, T., Hohsaka, T. & Matsuura, T. Different protein localizations on the inner and outer leaflet of cell-sized liposomes using cell-free protein synthesis. Synth. Biol. 3, (2018).

75. Schindelin, J. et al. Fiji: an open-source platform for biological-image analysis. Nat. Methods 9, 676–682 (2012).

76. Flory, P. J. & Rehner, J. Statistical Mechanics of Cross-Linked Polymer Networks II. Swelling. J. Chem. Phys. 11, 521–526 (1943).

77. Kang, M., Andreani, M. & Kenworthy, A. K. Validation of Normalizations, Scaling, and Photofading Corrections for FRAP Data Analysis. PLoS One 10, e0127966 (2015).

78. Soumpasis, D. M. Theoretical analysis of fluorescence photobleaching recovery experiments. Biophys. J. 41, 95–97 (1983).

79. Ganar, K. A. et al. Phase separation and ageing of glycine-rich protein from tick adhesive. Nat. Chem. 17, 186–197 (2025).

