## Supplementary Information for "Cyclical content exchange in hydrogel-core synthetic cells"

- **Supplementary Methods**
- **Supplementary Note 1**
- **Supplementary Figures**
    - Fig. S1-S16
- **Supplementary Tables**
    - Table S1-S6
- **Supplementary References**
- **Supplementary Movies**
    - Movies S1-S28

### Supplementary Methods

#### Cell-free expression of GFP in gel-core and liquid-core synthetic cells

Synthetic cells were prepared using the GUV phase transfer method as detailed in the **Methods**. The inner solution contained an adjusted buffer in order to balance the inner solution and additional TXTL components with the outer solution. Therefore, the inner solution contained the adjusted buffer (9.5 mM HEPES potassium salt, 120 mM potassium glutamate, 8.5 mM magnesium acetate, 58 mM glucose, 135 mM sucrose, 5 % w/v AF647-labelled $(PEG\text{-}SH)_4$, 5 mM glycy-l-tyrosine, pH 7.6) and in addition, the linear PCR product encoding GFP (prepared by amplifying the plasmid encoding eGFP; 5.9 ng $\mu L^{-1}$), 0.5 mM cysteine, 4 mM reduced glutathione (GSH) and PUREfrex 2.1 (GeneFrontier, Japan) Solution I, II and III at 4 µL, 0.5 µL and 1 µL per 20 µL of inner solution, respectively. For gel-core synthetic cells, 1 U $\mu L^{-1}$ HRP was included whilst for liquid-core synthetic cells, the equivalent volume of protein storage buffer (20mM HEPES, 250 mM potassium glutamate (pH 7.6)) was added.

Time-lapse movies were recorded on a Nikon Eclipse Ti2-U inverted microscope with CoolLED pE-300$^{white}$ illumination and a Nikon DS-Qi2 camera. Fluorescent probes were detected with FITC-3540C (GFP; Ex 475-nm; 4 ms exposure time (ET)), TRITC-B (rhodamine; Ex 550-nm; 226 ms ET) and Cy5-5070A (AF647; Ex 621-nm; 121 ms ET) filters and time-lapse imaging performed for 9.5 h with a 3 min interval (Plan Fluor ELWD 20x Ph1 ADM objective; final image pixel size: 0.33 µm/pixel).

#### The effect of the hydrogel on the activity of β-galactosidase

Samples were prepared in the wells of a 96-well plate. Firstly, 5 % w/v $(PEG\text{-}SH)_4$, 5 mM glycy-l-tyrosine and β-galactosidase (β-gal; 0, 0.1, 1.0, 10.0 U $\mu L^{-1}$) were mixed in 200 µL of buffer containing 20 mM HEPES potassium salt (pH 7.6) and 250 mM potassium glutamate. For gel samples, 1 U $\mu L^{-1}$ HRP was then added and mixed by pipetting up and down. For liquid samples, an equivalent volume of protein storage buffer (20mM HEPES/250 mM potassium glutamate) was added to the sample wells instead of HRP. After incubating for 2 hr to allow gelation, FDG (0.75 mM, dissolved in 20mM HEPES, 250 mM potassium glutamate (pH 7.6)) was then added to each well and the fluorescence intensity taken over time at 37°C using a plate-reader. Fluorescein was detected using excitation at 476-490 nm and emission collected from 515-545 nm. The gain was set at 200, the well-plate shaken at 200 rpm and the intensity measured every 3 min for 2 hr. 3 independent samples were collected for each condition (in 3 separate wells).

### Microscope Experimental Settings

#### Mal-PE binding in liquid-core synthetic cells

LSM 900; scanning zoom: 1x; pixel dwell time: 2.55 μs; pinhole: 1.00 A.U.; averaging: 16 (frame); final image pixel size: 0.20 μm/pixel.

#### Confirming the gelation of the hydrogel in the synthetic cell lumen

LSM 900 with AiryScan; scanning zoom: 1x; pixel dwell time: 1.03 μs; pinhole: 1.00 A.U.; averaging: none; final image pixel size: 0.31 μm/pixel.

#### Synthetic cell permeability at varying osmolarities, PEG concentrations and probe sizes

LSM 900; scanning zoom: 0.5x; pixel dwell time: 2.55 μs; pinhole: 1.34 A.U.; averaging: 2 (frame); final image pixel size: 0.40 μm/pixel.

#### Calcein release and subsequent encapsulation

LSM 900; scanning zoom: 1x; pixel dwell time: 0.76 μs; pinhole: 1.56 A.U.; averaging: none; final image pixel size: 0.096 μm/pixel.

#### Assessment of membrane fluidity and molecule diffusion in the gel network using FRAP

LSM-900; images were captured sequentially with 2.5 x scanning zoom, a pixel dwell time of 1.00 μs and a 1.63 AU pinhole and no averaging. The final image pixel size was 0.08 μm/pixel (NBD-PE) or 0.13 μm/pixel (FITC-dextran).

#### Enzymatic reaction in the synthetic cell lumen using substrates supplied from outside

LSM 900; scanning zoom: 0.5x; pixel dwell time: 2.55 μs; pinhole: 1.56 A.U.; averaging: 2 (frame); final image pixel size: 0.40 μm/pixel.

**Supplementary Note 1**

<u>Theoretical estimation of the gel mesh size</u>

The mesh size, ξ, can be approximated from the following modified form of the Canal-Peppas equation,[1] as proposed by Richbourg et al.: [2]

$$\xi = \varphi_s^{-1/3}[\left(1-\frac{2}{f}\right)\bar{l}^2 C_\infty \lambda N_j]^{1/2} \qquad \text{Eq. S1}$$

Where $\varphi_s$ is the swollen polymer volume fraction, *f* is the junction functionality, $\bar{l}$ is the average bond length in the polymer repeat unit, $C_\infty$ the Flory characteristic ratio of the polymer (4 for PEG; this is a measure of the stiffness of the polymer), $\lambda$ the number of backbone bonds per repeat unit (3 for PEG), and $N_j$ the degree of polymerization between junctions. Since each PEG arm is ~5 kDa in size, corresponding to 112 PEG units, for two arms joined between each junction there are ~224 monomer units.

For a hydrogel network composed of 20 kDa 4-arm thiolated PEG, each repeat PEG unit has 2 × C-O and 1 × C-C bond. Given an average C-O bond length of 0.143 nm and C-C bond length of 0.154 nm, this gives an average bond length in PEG of ((2 × 0.143)+0.154)/3) = 0.15 nm.

Therefore, for the gel-core synthetic cells encapsulating 4-arm PEG at 2.5, 5.0 and 7.5 % w/v, the mesh sizes, ξ, are calculated as 19.9, 15.8 and 13.8 nm respectively. The mesh radius $r_m$ can then be calculated given *f* = 4 as follows:

$$r_m = \frac{\sqrt{6}}{3}\xi \qquad \text{Eq. S2}$$

This gives mesh radii of 16.2, 12.8 and 11.2 nm respectively for [PEG-$SH_4$] at 2.5, 5.0 and 7.5 % w/v.

### Supplementary Figures

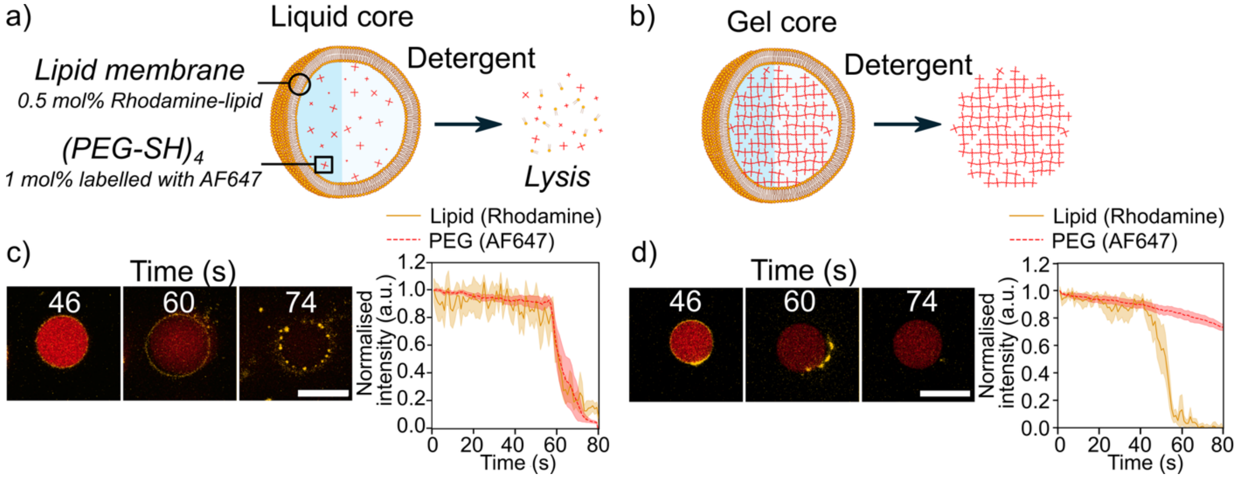


**Figure S1** – **The effect of detergent on the synthetic cells.** Schematic showing response of rhodamine lipid-labelled synthetic cells encapsulating AF647-PEG to detergent; liquid-core synthetic cells completely lysed (a), whereas gel-core synthetic cells underwent membrane lysis with retainment of the hydrogel core (b). (c,d) Confocal micrographs over time (left) and associated line plot of mean fluorescence intensity over time (right) showing the change in fluorescence signal of rhodamine-PE and AF647-PEG for liquid-core (c) and gel-core (d) synthetic cells. $n$=7 (liquid), 9 (gel); shaded regions in the line plots indicate ± 1 s.d. from the mean. The data were normalised by subtracting the background and scaling, with 1 being the maximum intensity. Detergent was added to the samples at 0 s. All scale bars=20 μm.

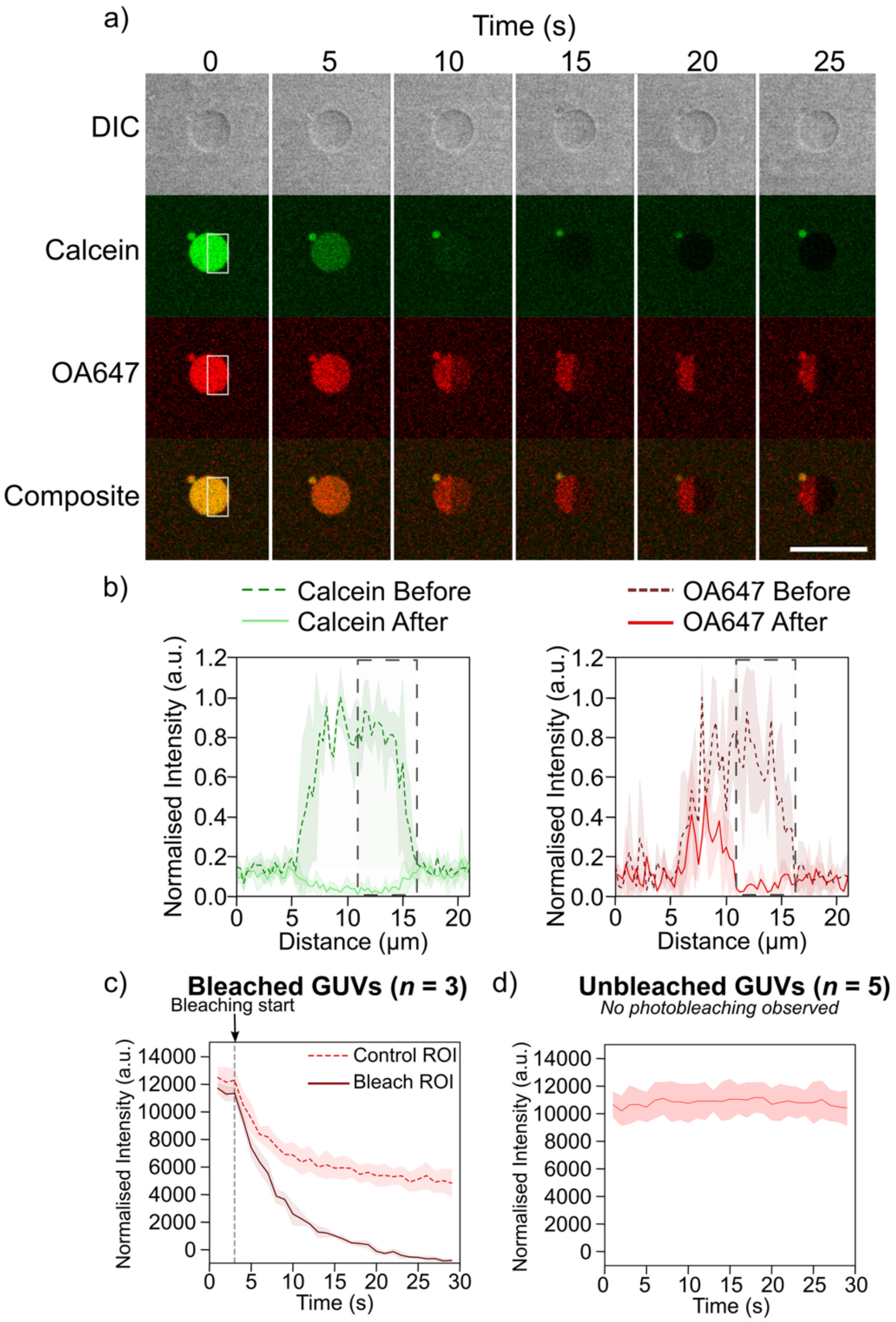


**Figure S2 – The crosslinked $(PEG\text{-}SH)_4$ network facilitates protein immobilization in gel-core synthetic cells.** (a) Confocal micrographs of a gel-core synthetic cell encapsulating calcein and OA647 before and after sequential photobleaching and image capture; the bleach ROI is indicated by the white box. Simultaneous photobleaching with the 488-nm and 647-nm lasers was conducted. Calcein fluorescence intensity rapidly diminished throughout the lumen, ultimately dropping below that of the outside solution due to the cumulative effects of repeated photobleaching. On the other hand, OA647 fluorescence reduced dramatically in the bleach ROI but to a lesser degree in the unbleached region, indicating immobilization of the OA647 protein. Scale bar=20 µm. (b) Mean plot profile of gel-core synthetic cells of similar diameter to that in (a) encapsulating calcein (left) and OA647 (right) before (0 s) and after (25 s) photobleaching (*n*=3). The location of the bleach ROI is indicated as the dashed box. The data were normalised by subtracting the background and scaling, with 1 being the maximum intensity. The plots are consistent with our finding that proteins are immobilised in the gel network. (c) Mean fluorescence profiles of OA647 intensity over time in synthetic cells within an unbleached control ROI and within a bleach ROI (647-nm laser). Bleach ROIs are as shown in (a), covering approximately half of the synthetic cell equatorial area, whilst control ROIs cover the remaining equatorial area. Photobleaching commenced after 3 s and continued for 26 s with an image collected every second (*n*=3). The decrease in fluorescence in the control ROI is due to unbound OA647 diffusing from the control ROI into the bleach ROI. Once all the unbound OA647 had been photobleached, the bleach ROI fluorescence intensity decreased to zero, with the residual fluorescence intensity in the control ROI corresponding to immobilised OA647 that was crosslinked to the gel-core, and which could not enter the photobleaching ROI. (d) Fluorescence plot of OA647 intensity over time in the lumen of synthetic cells that were not explicitly photobleached indicates that there is minimal photobleaching over the course of the image acquisition (*n*=5). Intensity values in (c-d) were normalised by subtracting the background signal for each synthetic cell before averaging. Shaded regions in the line plots indicate the mean ±1 s.d.

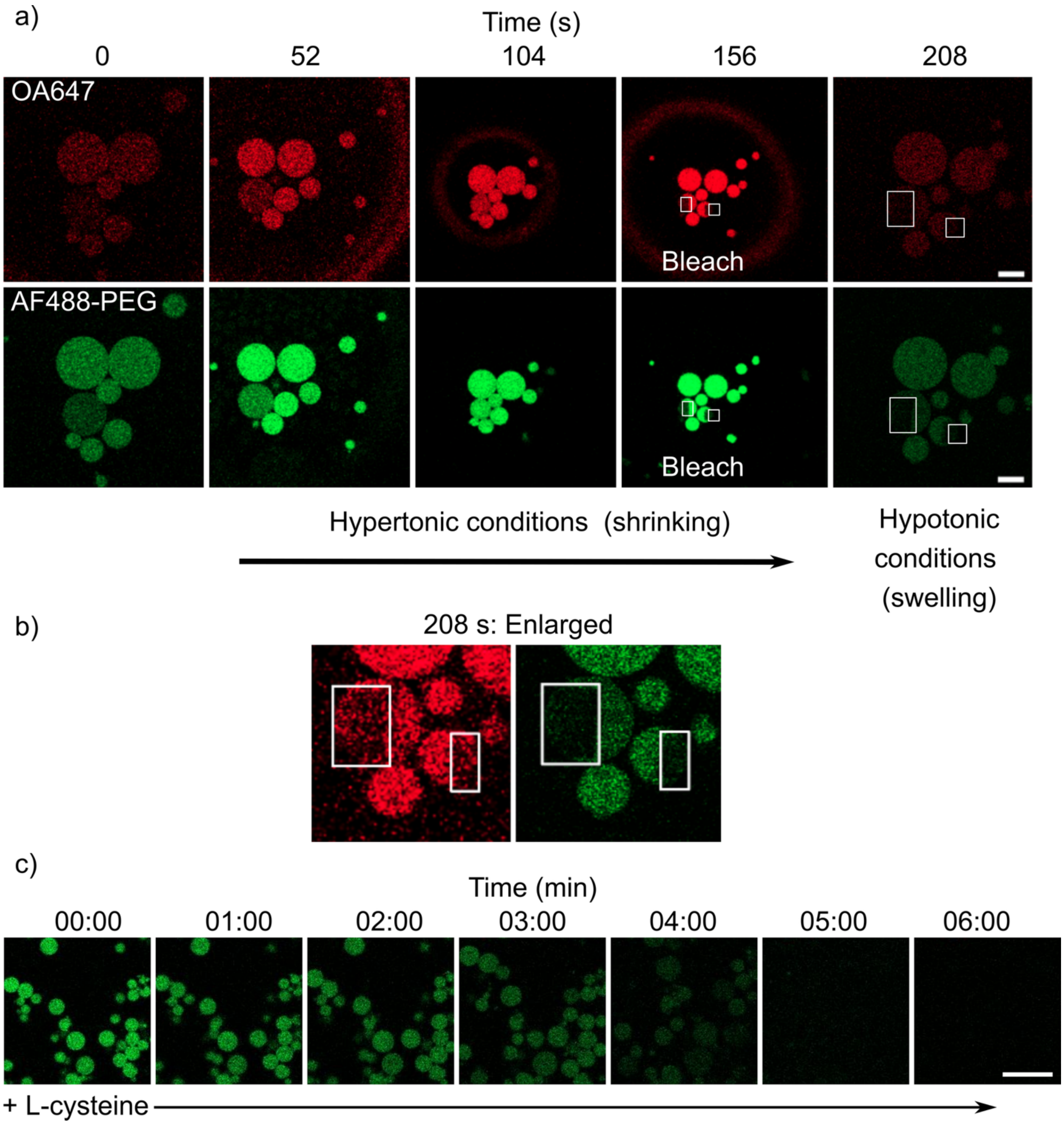


**Figure S3 – Characterization of membraneless hydrogels.** Membraneless gels were prepared by detergent-induced membrane lysis of gel-core synthetic cells to leave behind the uncoated gel cores. The gel core was labelled with AF488, whilst OA647 was immobilised in the core via cysteine-thiol conjugation. (a) The membraneless gels showed shrinking in size under hypertonic conditions and swelling under hypotonic conditions, with an associated increase and reduction in fluorescence due to compression (0 to 156 s) and expansion (156 to 208 s) in size. The buffer conditions are the same as for gel-core synthetic cells in Fig. 3. After shrinking the gels, two ROIs of OA647 or AF488-PEG (white boxes) were photobleached simultaneously using the 647-nm or 488-nm laser lines, respectively. Both fluorophores photobleached within the ROI and did not show recovery, indicating immobilization of both AF488 and OA647 (156 s). These photobleached ROIs remained in the swollen gel and did not recover over time, indicating that the gels retained their structure even after swelling (208 s). (b) Enlarged view of the ROIs from the rightmost panel of (a); contrast was adjusted for visualization. (c) Confocal micrographs over time showing the degradation of membraneless gel cores after the addition of 1 mM L-cysteine (Cys) (at 0 s). The AF488-labelled gel cores were produced after detergent-induced lysis of the lipid membrane of gel-core synthetic cells, and Cys was added in buffer isotonic with the synthetic cell solution. The gels first expanded before completely breaking apart. This indicates that the hydrogel was disassembled into the constituent $(PEG\text{-}SH)_4$ units. The images also indicate that the rate-limiting step in gel disassembly is the reduction of the disulfide bonds by Cys, as opposed to the diffusion of Cys into the gel network. If diffusion of Cys were to be the rate-limiting step, the disassembly of the gel core would be expected to proceed from the outside-in, with the gels becoming smaller over time, while we observed a homogeneous decrease in the fluorescence regardless of the location. All scale bars=20 µm.

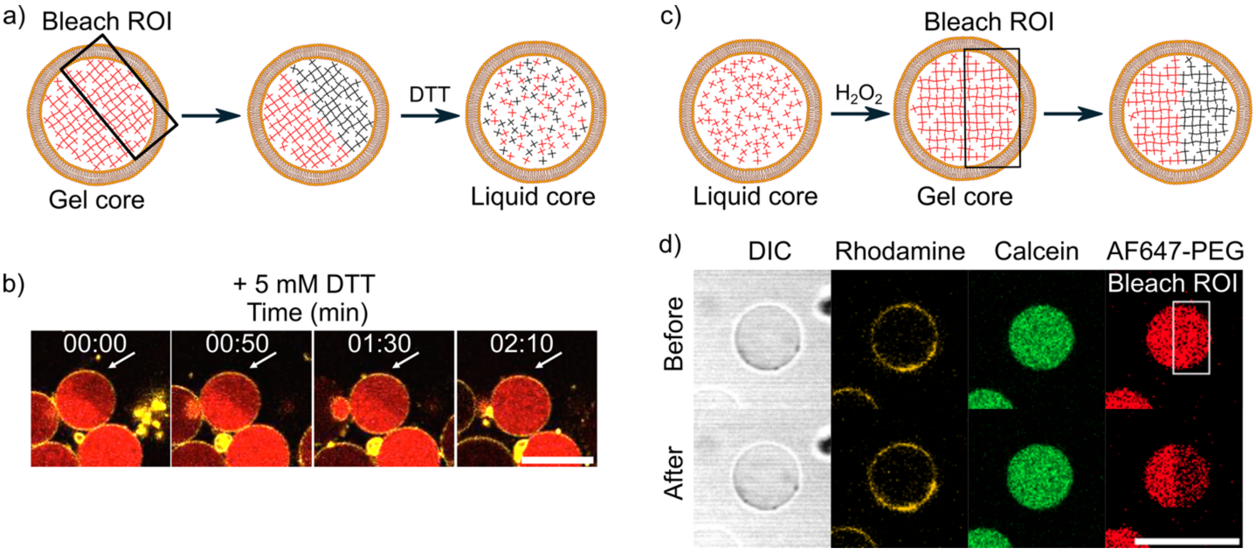


**Figure S4 – The gel network inside synthetic cells can be disassembled and reassembled by the addition of redox agents.** Rhodamine lipid-labelled synthetic cells encapsulating AF647-PEG are shown. (a) Schematic and (b) confocal micrographs over time showing the reduction of disulfide bonds in the AF647-labelled gel core, causing gel disassembly to a liquid core with homogenization of the photobleached ROI (white arrows) after addition of 5 mM DTT to the sample surface, at the top of the imaging chamber. The time indicates the time that DTT was added, and therefore the delay between DTT addition and gel disassembly is partly due to the time taken for macroscopic diffusion of DTT from the sample surface to the synthetic cell location. (c) Schematic showing synthetic cells with a liquid core that resulted from gel-core disassembly using DTT (left), which were then reoxidised to a gel network using 10 mM $H_2O_2$ (middle), before photobleaching to verify the state of the core (right). (d) Confocal micrographs of synthetic with a gel core that had been disassembled and then reassembled to a gel network using 10 mM $H_2O_2$ (top). The same synthetic cell is shown before (top) and after (bottom) photobleaching of the AF647, with the white box indicating the bleach ROI. Encapsulated calcein was not released, indicating maintenance of the membrane barrier function throughout gel disassembly and reassembly. All scale bars=20 µm.

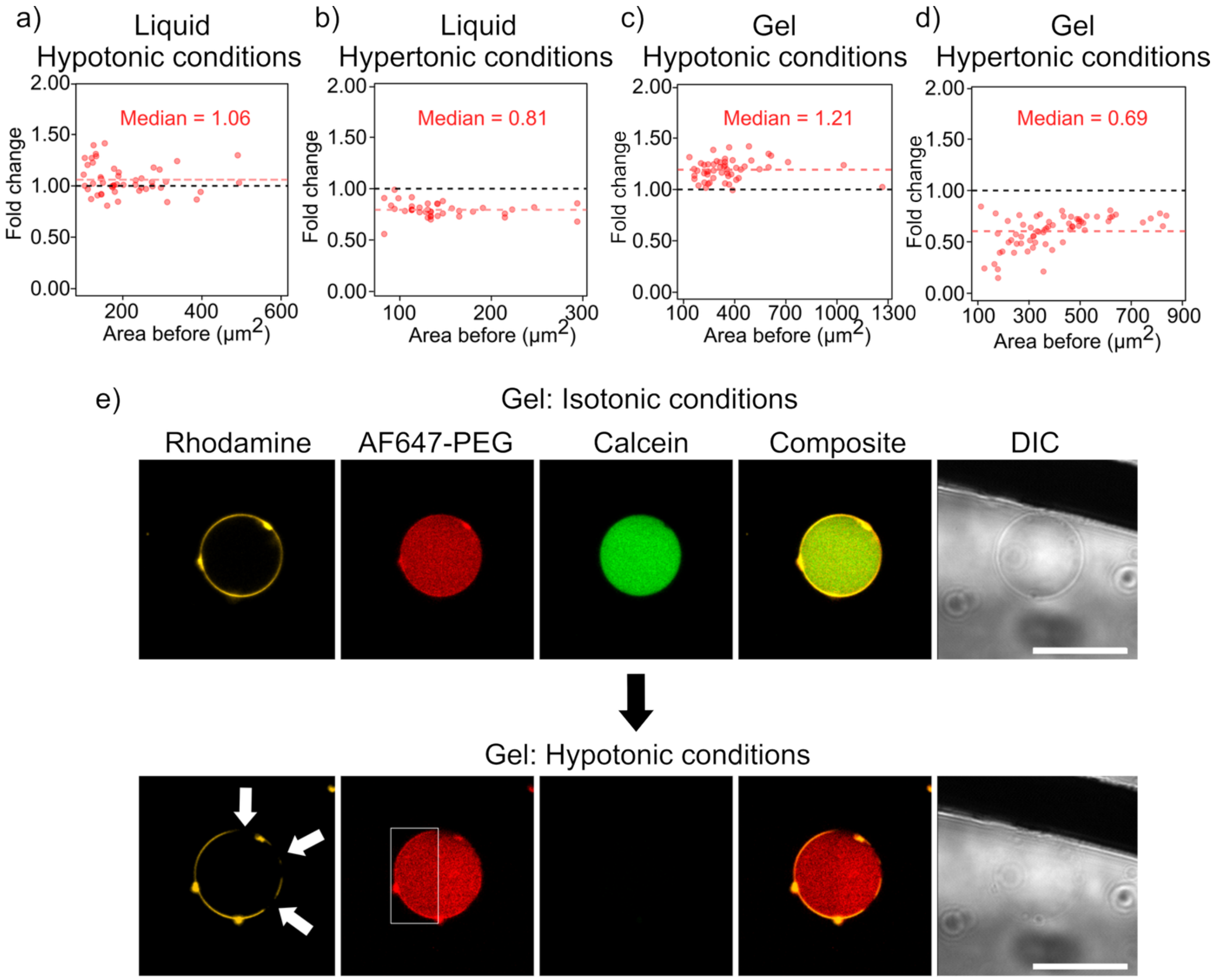


**Figure S5 – Swelling and shrinking of synthetic cells**. Rhodamine lipid-labelled synthetic cells encapsulating AF647-PEG and calcein are shown. (a-d) Plots indicating the fold change in size of synthetic cell cross-sectional areas before and after hypotonic or hypertonic exposure for liquid-core (a, b) and gel-core (c, d) synthetic cells, using the same dataset and cross-sectional area as defined in Figs. 2c,d. From the median fold change, it can be seen that the size change is larger for gel-core synthetic cells and there is no indication of dependence on initial size (*n*=49, 39 (liquid-core: left, right), *n*=66, 52 (gel-core: left, right)). The dashed black line at *y*=1 indicates no change in size; the pink dashed line indicates the median value. (e) Representative confocal micrograph of a gel-core synthetic cell under isotonic conditions (top) and after exposure to hypotonic conditions (bottom). Hypotonic exposure leads to gel swelling and membrane defect formation visible as dark regions in the membrane (rhodamine channel) and loss of calcein due to permeation through the membrane defects. The gel increases in size, as visible from the rhodamine channel. Photobleaching of the gel is visible in the left-hand side of the synthetic cell in the AF647-PEG channel after swelling (white box), indicating maintenance of a gel network. The loss in density difference between the synthetic cell interior and exterior due to content exchange through the membrane defects also leads to the synthetic cell becoming less visible on the DIC channel (rightmost panel). All scale bars=20 μm.

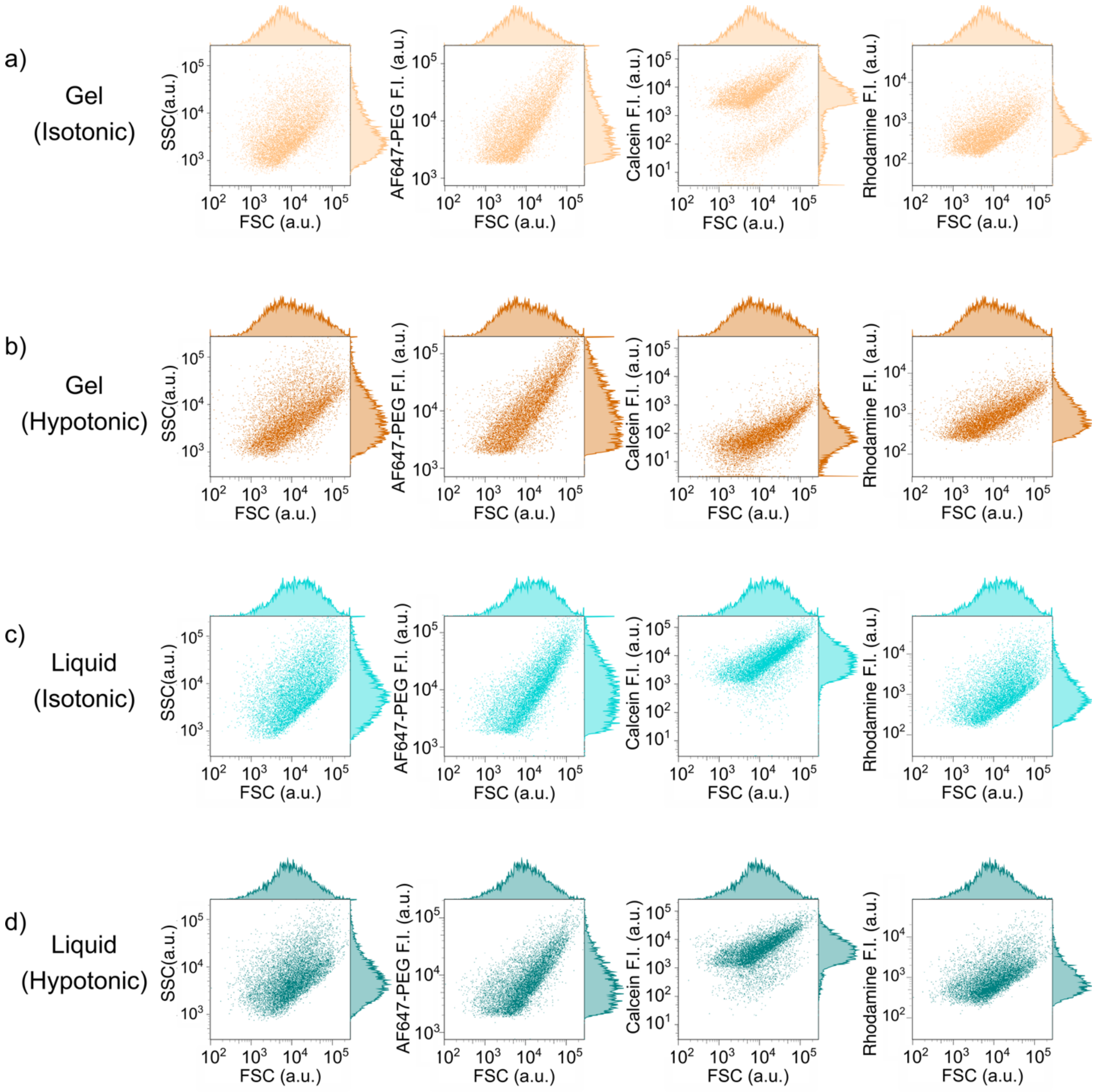


**Figure S6 – Extended data of Fig. 2f,h.** Two-dimensional FC dot plots for gel-core (a, b) and liquid-core (c, d) synthetic cells without mal-PE in the membrane under isotonic and hypotonic (swelling) conditions. The leftmost plots in each condition show forward scattering (FSC) versus side scattering (SSC), and remaining plots show FSC versus each fluorophore in the synthetic cell, with the fluorophore signal corresponding to fluorescence intensity. AF647-PEG indicates the PEG monomer (liquid-core synthetic cells) or gelled network (gel-core synthetic cells), calcein (10 μM) is encapsulated in the lumen and rhodamine (1 mol%) in the membrane. *n*=10000 for all.

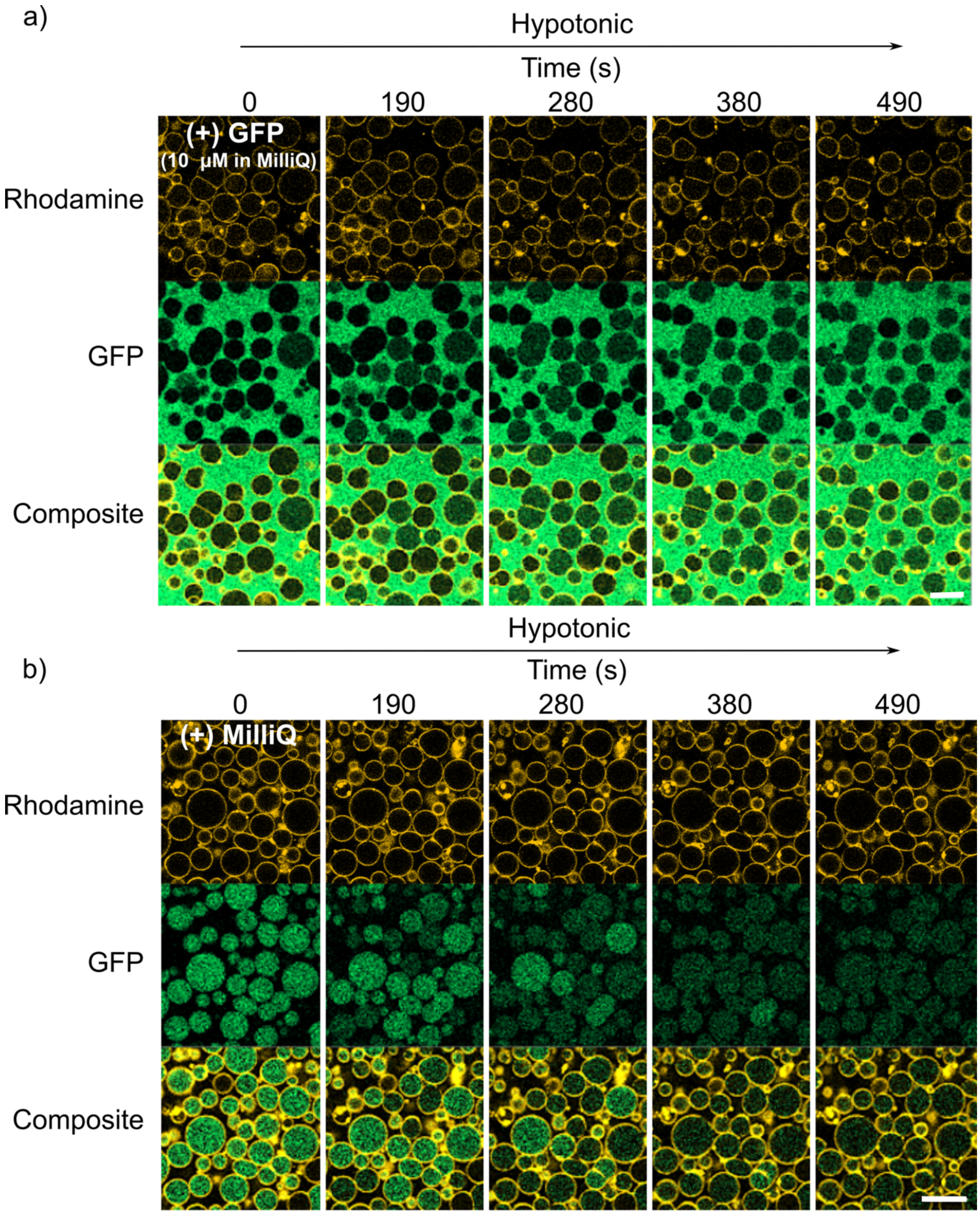


**Figure S7 – Extended data of Fig. 3c: Gel-core synthetic cells swell to capture or release GFP.** (a) Confocal micrographs indicating the influx of GFP into gel-core synthetic cells that initially did not encapsulate GFP. The synthetic cells swell in hypotonic conditions, facilitating the permeation of GFP from the external solution through membrane fissures. (b) Confocal micrographs indicating the efflux of GFP from gel-core synthetic cells when exposed to hypotonic conditions. The synthetic cells swell, leading to release of the mobile GFP from inside the synthetic cells through induced membrane fissures. A portion of GFP was seen to be immobilised inside the lumen, due to disulfide crosslinking between available cysteine residues on GFP and free thiols in the gel network. All scale bars=20 µm.

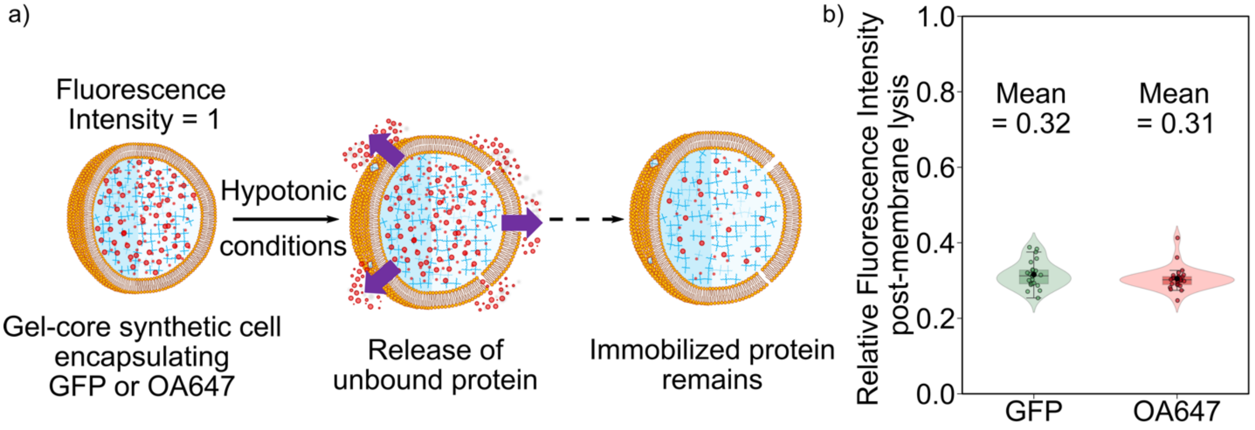


**Figure S8** – **A portion of encapsulated proteins remain immobilised in gel-core synthetic cells after swelling.** (a) Schematic showing the swelling of gel-core synthetic cells encapsulating either GFP or OA647 in hypotonic conditions, with release of any unbound protein via induced membrane fissures. (b) Violin plot indicating the relative fluorescence intensity inside gel-core synthetic cells encapsulating 10 µM of either GFP (left, green) or OA647 (right, red) after hypotonic swelling as shown in (a). The initial fluorescence intensity before swelling was set to 1. Each coloured circle indicates one synthetic cell datapoint (black circles indicate the mean). Both systems swell in hypotonic buffer to induce membrane fissure formation and release of any unbound protein. Approximately 70% of encapsulated protein was released in both systems, with the final fluorescence intensity for both fluorescent proteins equal to roughly one third of the initial intensity ($n$=23 (GFP): $n$=26 (OA647). The GFP data is the same as used in Figure 3c (GFP-efflux).

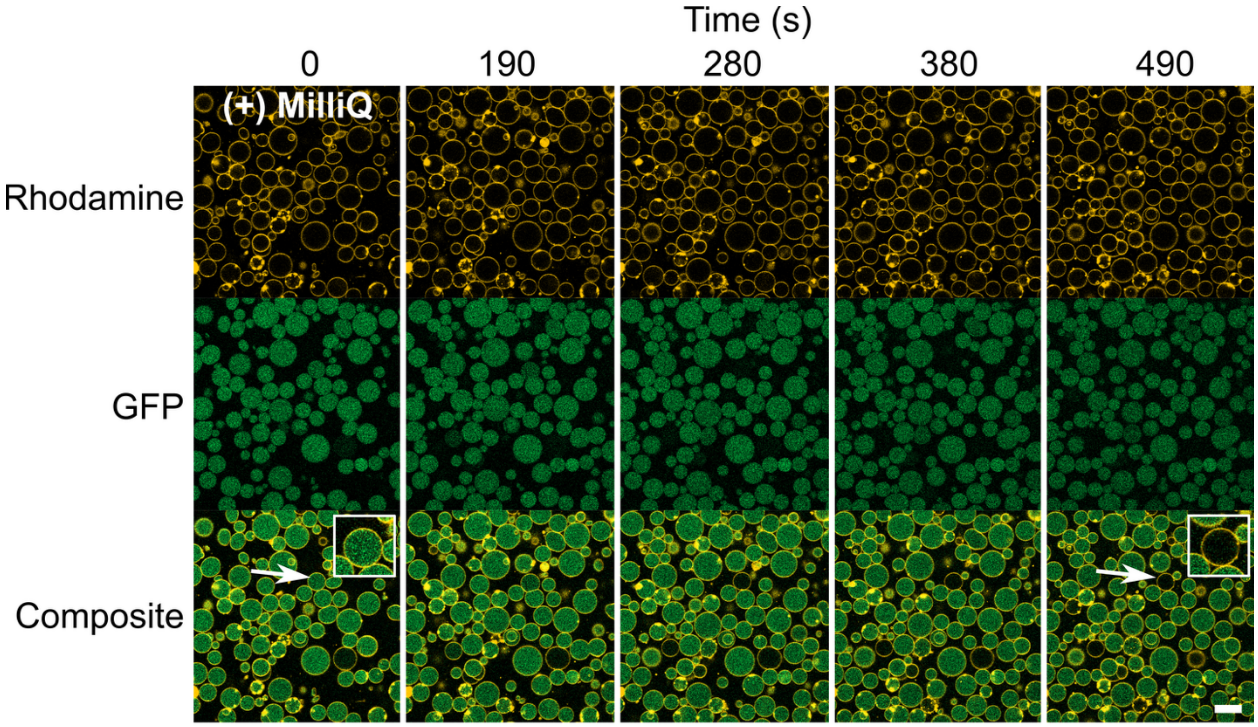


**Figure S9 – Extended data of Fig. 3c: Liquid-core synthetic cells do not swell to release GFP.** (a) Confocal micrographs indicating minimal efflux of GFP from liquid-core synthetic cells when exposed to hypotonic conditions. Unlike gel-core synthetic cells (Fig. S7), the majority of liquid-core synthetic cells do not swell sufficiently in hypotonic conditions to release GFP through membrane fissures. However, one exemplar synthetic cell is shown in the inset (bottom row, white arrow) that did show release, presumably via transient pore formation in the membrane as the osmolarity equilibrated between the lumen and external buffer. All scale bars=20 μm.

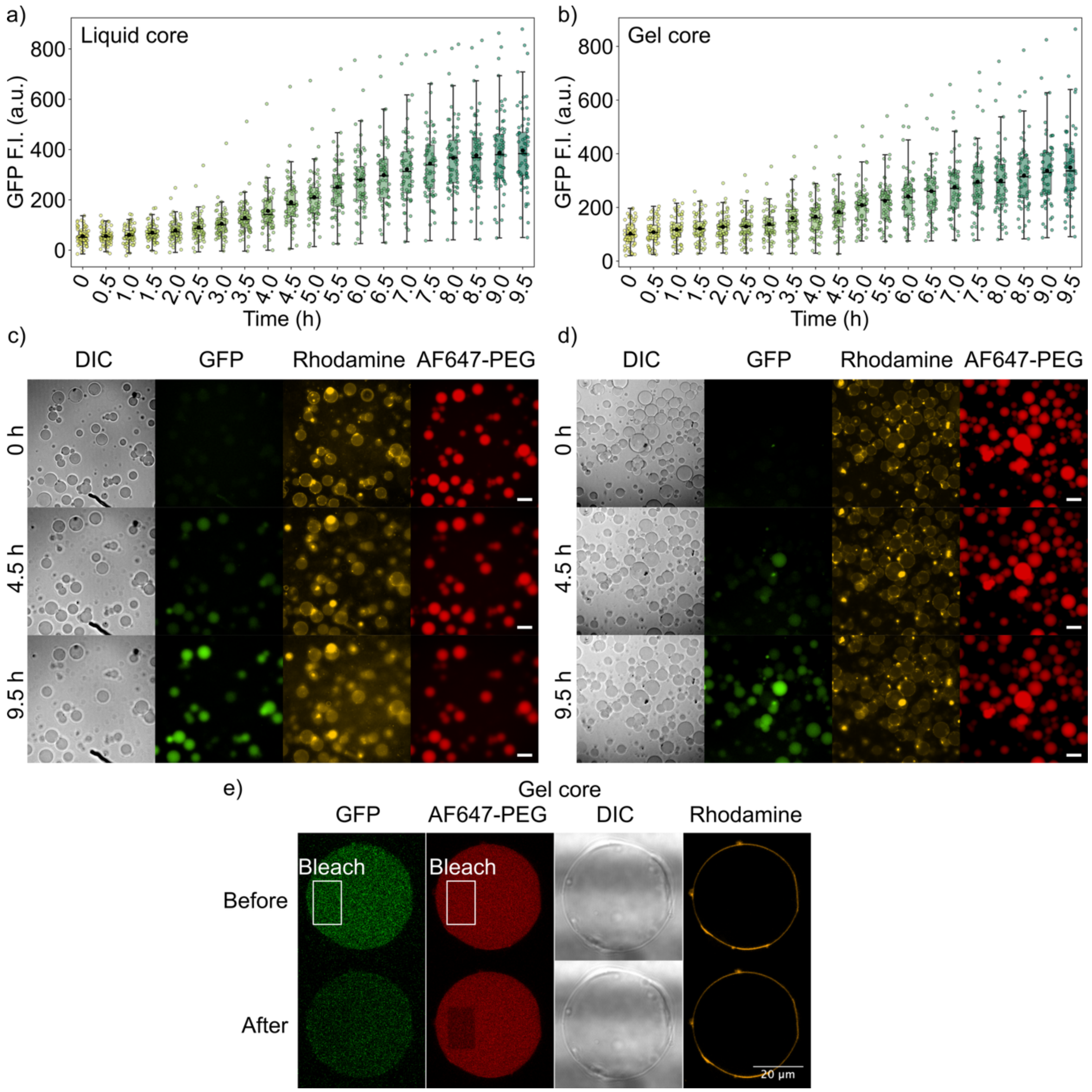


**Figure S10** – **Expression of GFP in liquid- and gel-core synthetic cells using the PUREfrex system.** (a-b) Violin plots indicating the GFP fluorescence intensity over time in liquid-core (a, *n*=68) and gel-core (b, *n*=63) synthetic cells. The box-plot centre lines depict the median and the boxes span the interquartile range (25–75%); the black circles indicate the mean at each time-point. (c-d) Confocal micrographs corresponding to some of the datapoints in (a-b) indicating the expression of GFP (green) over time in synthetic cells with a liquid core (c) or gel core (d). The membrane contains rhodamine-PE (yellow) and the encapsulated $(PEG\text{-}SH)_4$ is labelled with AF647 (red). (e) Confocal micrographs indicating the fluorescence in a GFP-expressing gel-core synthetic cell before (top) and after (bottom) photobleaching, performed 10 hours after expression of GFP was initiated. The GFP and AF647 fluorophores were photobleached at 100% for 100 iterations in the indicated rectangular ROI (white boxes). The AF647-PEG only photobleached in the photobleaching ROI, indicating that the lumen is in a gel state, while the GFP fluorescence reduced throughout the lumen, indicating that the GFP remained mobile. This is likely due to the presence of DTT and Cys in the PUREfrex system. Hence whilst GFP can be coupled to the hydrogel via disulfide bridges, the presence of the reducing reagents may have reduced the coupling efficiency in this case. All scale bars=20 µm.

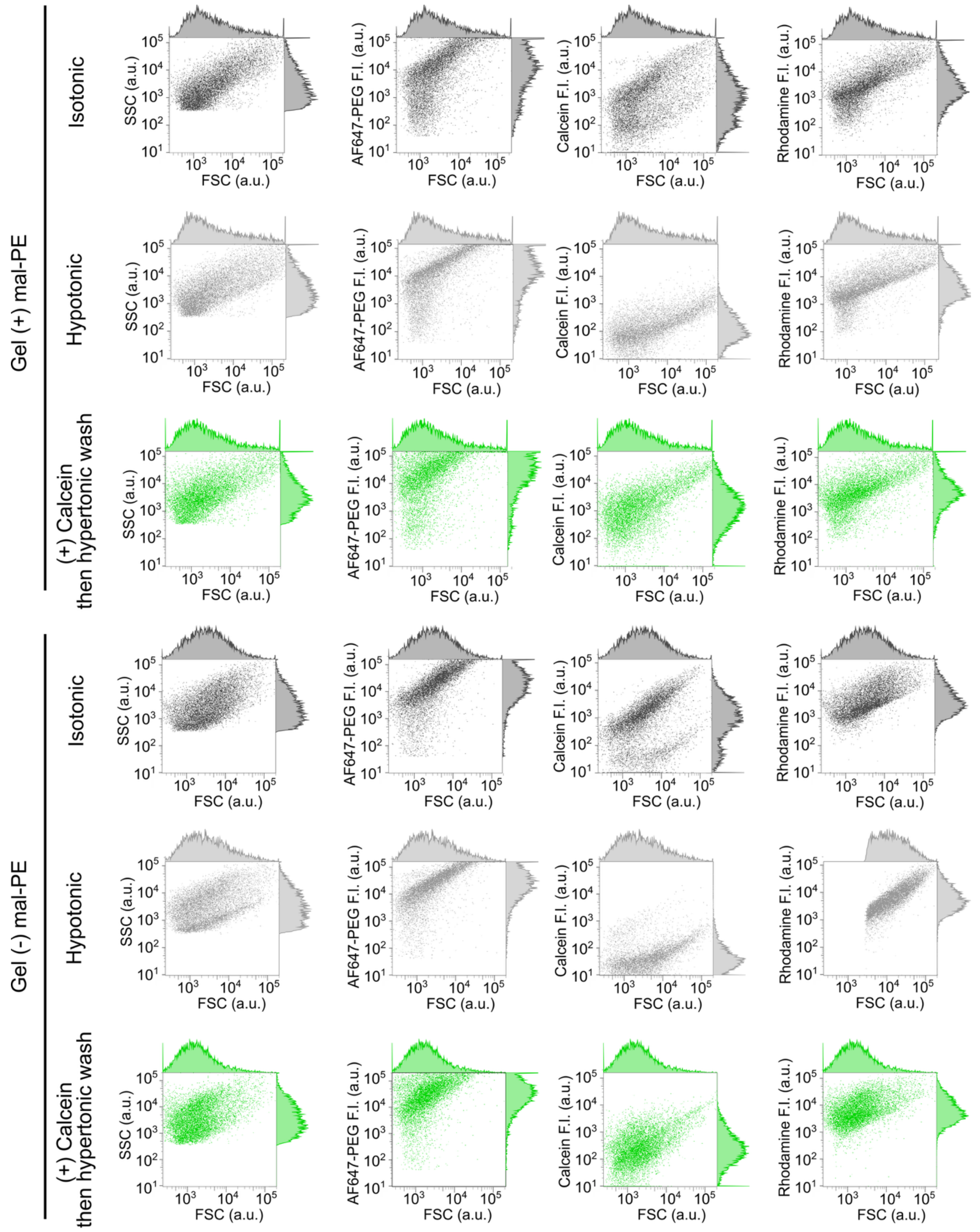


**Figure S11** – **Extended data of Fig. 4f.** (a) Two-dimensional FC dot plots for gel-core synthetic cells with and without mal-PE in the membrane under different conditions. *n*=10805, 10556, 10019, 10329, 10192, 10210 (top to bottom).

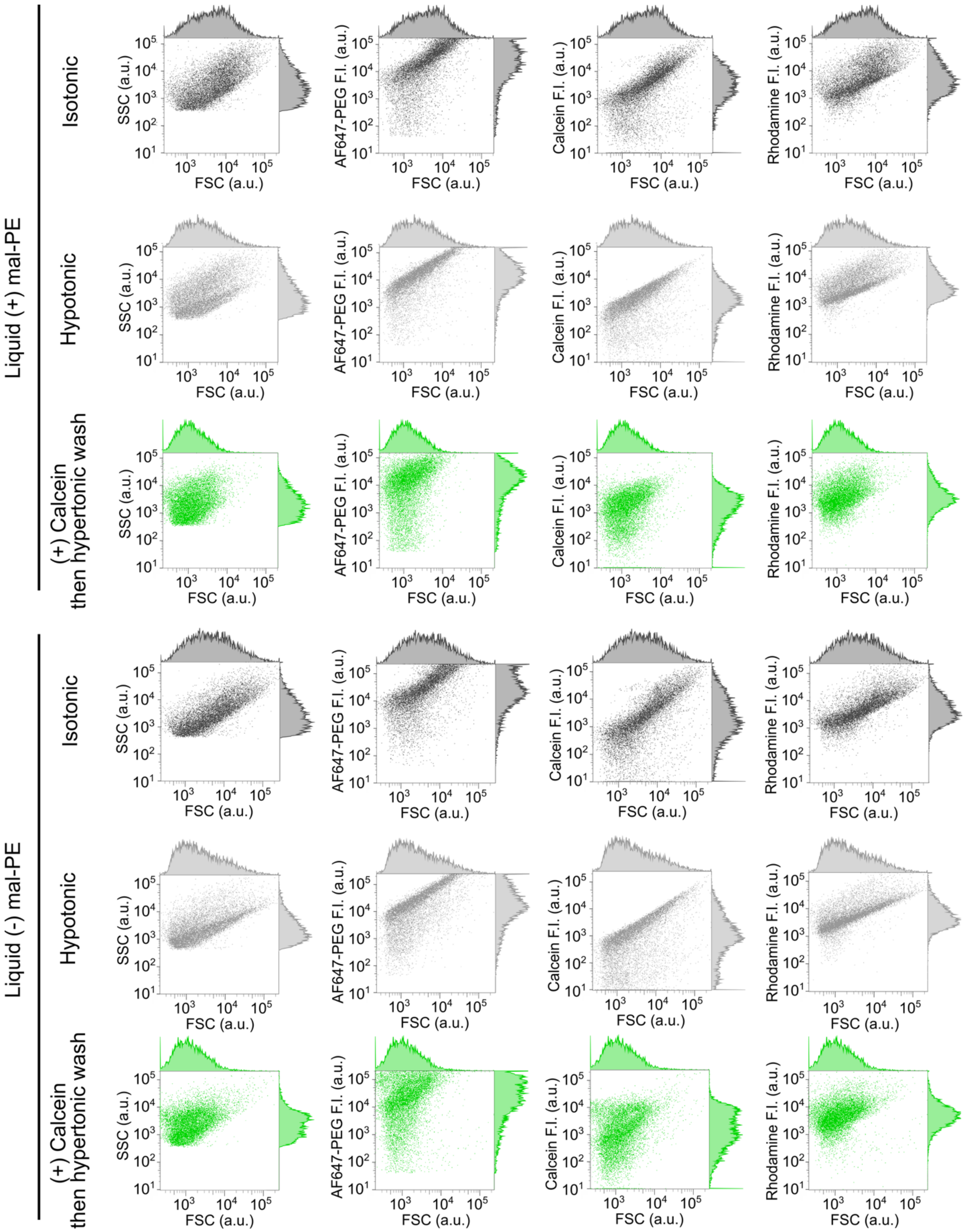


**Figure S12** – **Extended data of Fig. 4f.** (a) Two-dimensional FC dot plots for liquid-core synthetic cells with and without mal-PE in the membrane under different conditions. *n*=10186, 10339, 10726, 10497, 10258, 10873 (top to bottom).

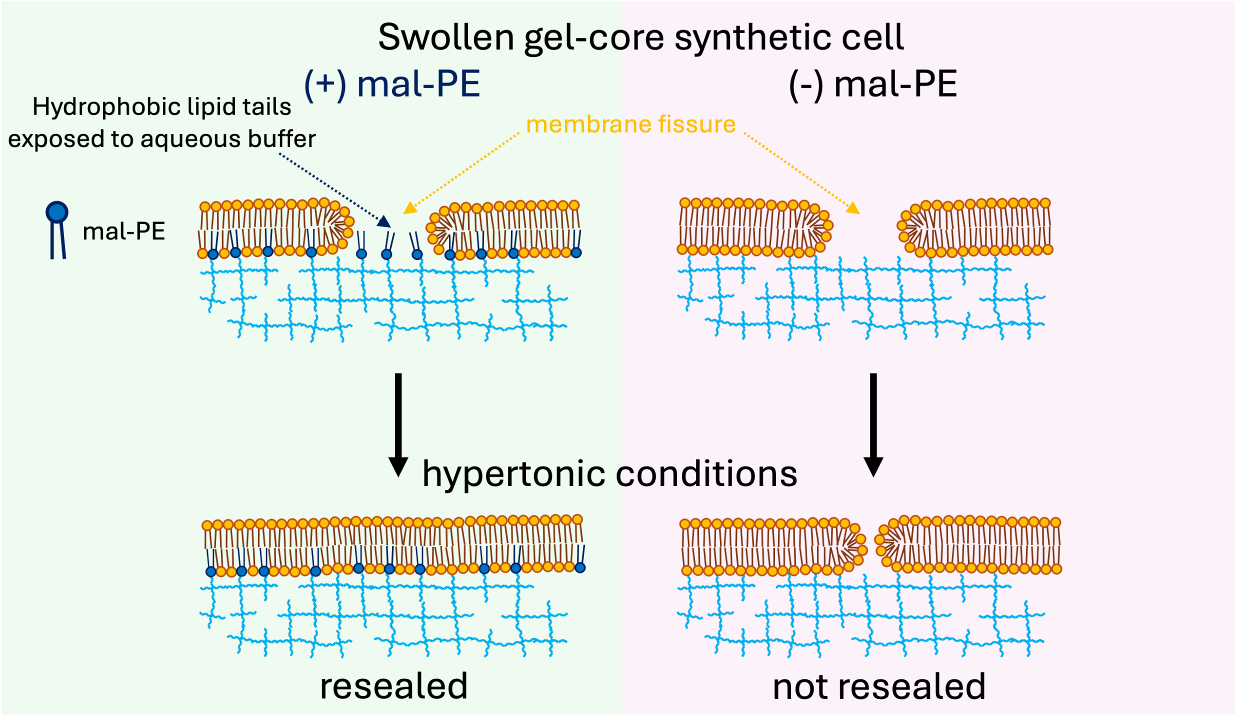


**Fig S13 – Schematic of membrane resealing during the shrinking of swollen gel-core synthetic cells**. Proposed mechanism of the effect of mal-PE on membrane fissure resealing. The schematic indicates gel-core synthetic cells with membrane fissures formed after swelling in hypotonic conditions (top). Regardless of the presence or asbsence of mal-PE, the internal hydrogel shrinks under hypertonic conditions to lower the overall membrane surface area and bring the membrane edges into proximity (bottom schematic). For complete resealing of the membrane fissure, a fusion stalk must form between the apposing membrane edges. This would expose the hydrophobic acyl chains of the lipids on the membrane edges to aqueous buffer, presenting an energetic barrier to resealing in the absence of mal-PE (bottom right). In this condition, lipid aggregates are seen to form on the gel surface after gel shrinking, and resealing of the fissures is not seen (Fig. 4d). Conversely, when mal-PE is present, anchor points are formed in the membrane where mal-PE is crosslinked to the internal hydrogel. The hydrophobic acyl chains of these anchored mal-PE lipids are exposed to the aqueous solution during hypotonic swelling and membrane fissure formation, which is energetically unfavourable (top left). During shrinking of the gel, these anchored mal-PE lipids act as sites for membrane rewetting and reduce the energetic penalty of membrane resealing (bottom left). The key step for membrane resealing is this rewetting-mediated membrane fusion, enabled by the anchored mal-PE lipids.

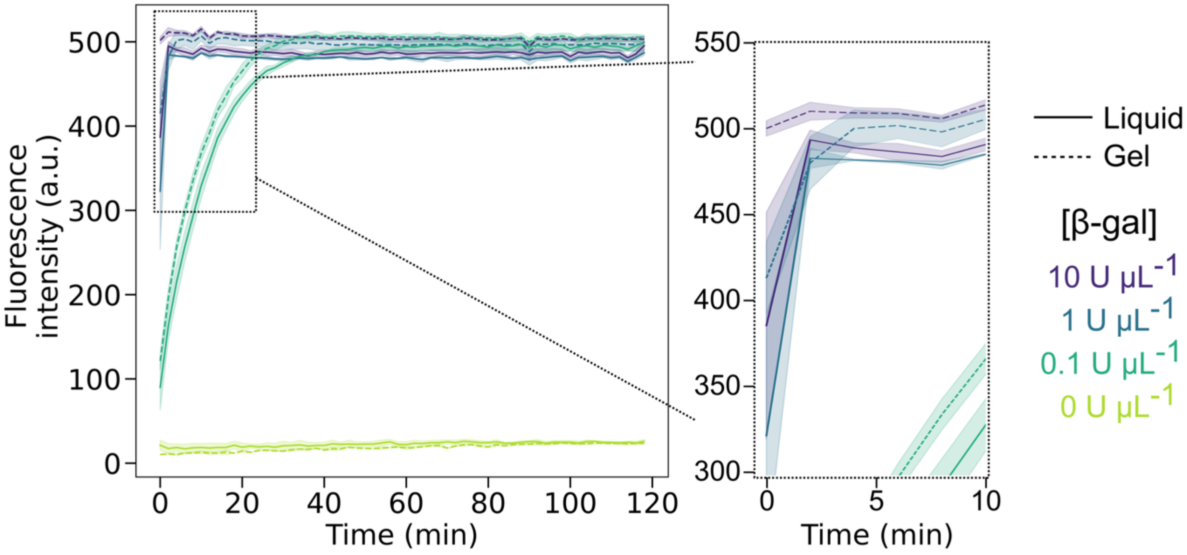


**Fig S14** – **Assessing the effect of the hydrogel on β-gal hydrolysis activity.** Line plots of the fluorescence intensity over time indicating the hydrolysis of FDG to fluorescein by β-gal in bulk liquid or gel samples, prepared in a 96-well plate (see **Supplementary Methods**). Lines and shaded region indicate the mean ±1 s.d. for each condition (*n*=3 for all). The presence of the gel (dashed lines) did not significantly influence the reaction rate, when compared with the equivalent reaction conditions in a liquid sample (solid lines. 1 U μL$^{-1}$ β-gal was sufficient for the reaction to reach a plateau within 10 min and therefore this concentration was used in the subsequent experiments in gel- and liquid-core synthetic cells.

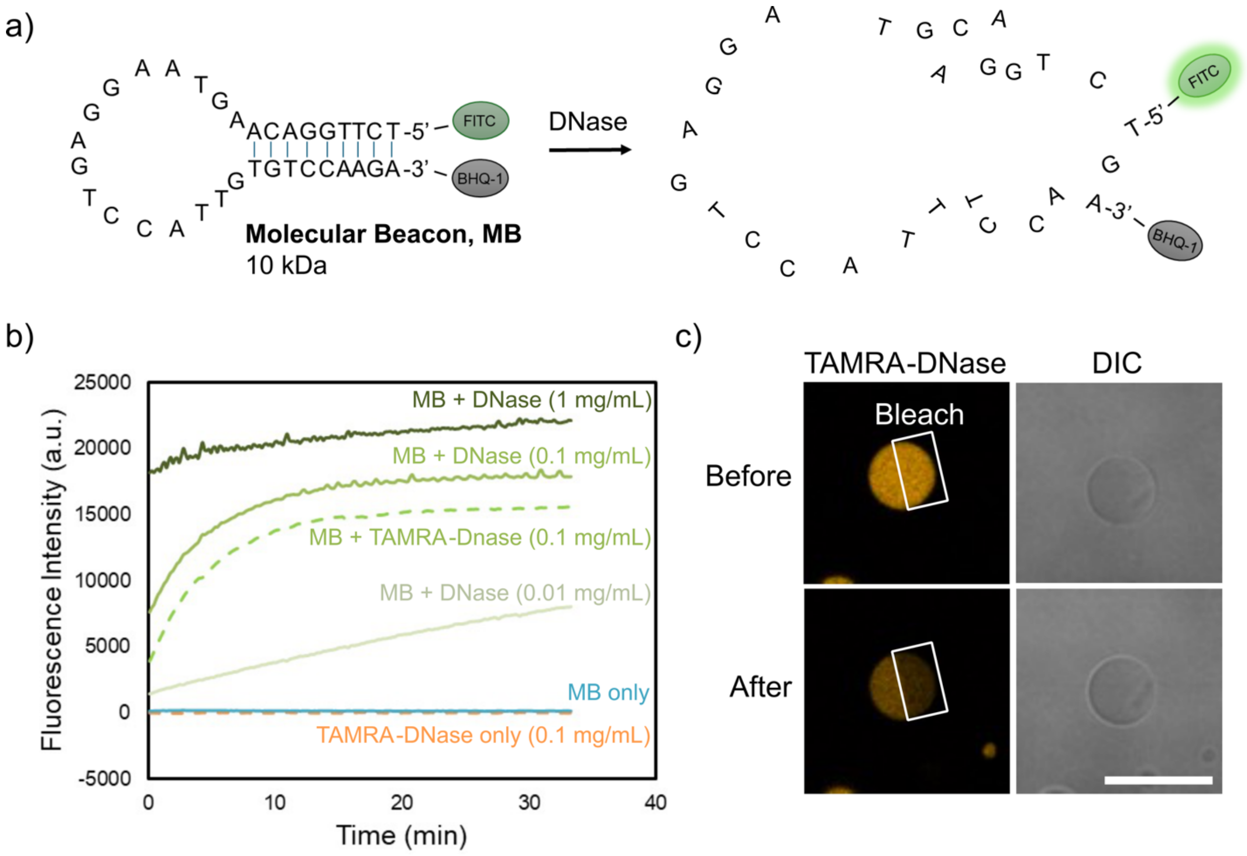


**Figure S15 – Molecular beacon degradation via DNase and TAMRA-DNase immobilization in gel-core synthetic cells.** (a) Schematic showing the molecular beacon- (MB-) DNA oligonucleotide sequence and structure, with fluorescence emitted upon MB-DNA digestion via DNase and unquenching of FITC. (b) Line plots indicating that FITC fluorescence increased over time when the MB-DNA is mixed with DNase at the concentrations indicated. In bulk conditions, DNase labeled with TAMRA (TAMRA-DNase) exhibited similar activity to unlabeled DNase. TAMRA-DNase was then used for visualization of the enzyme inside synthetic cells. (c) Confocal micrographs of gel-core synthetic cells with TAMRA-DNase immobilized on the hydrogel, before and after photobleaching (white box). The TAMRA fluorescence reduced to a larger extent in the bleach ROI, indicating immobilization of TAMRA-DNase. Some reduction in fluorescence was also seen in the non-photobleached region, indicating a mobile fraction of TAMRA-DNase (as noted for other proteins; **Fig. S2**). Scale bar=20 μm.

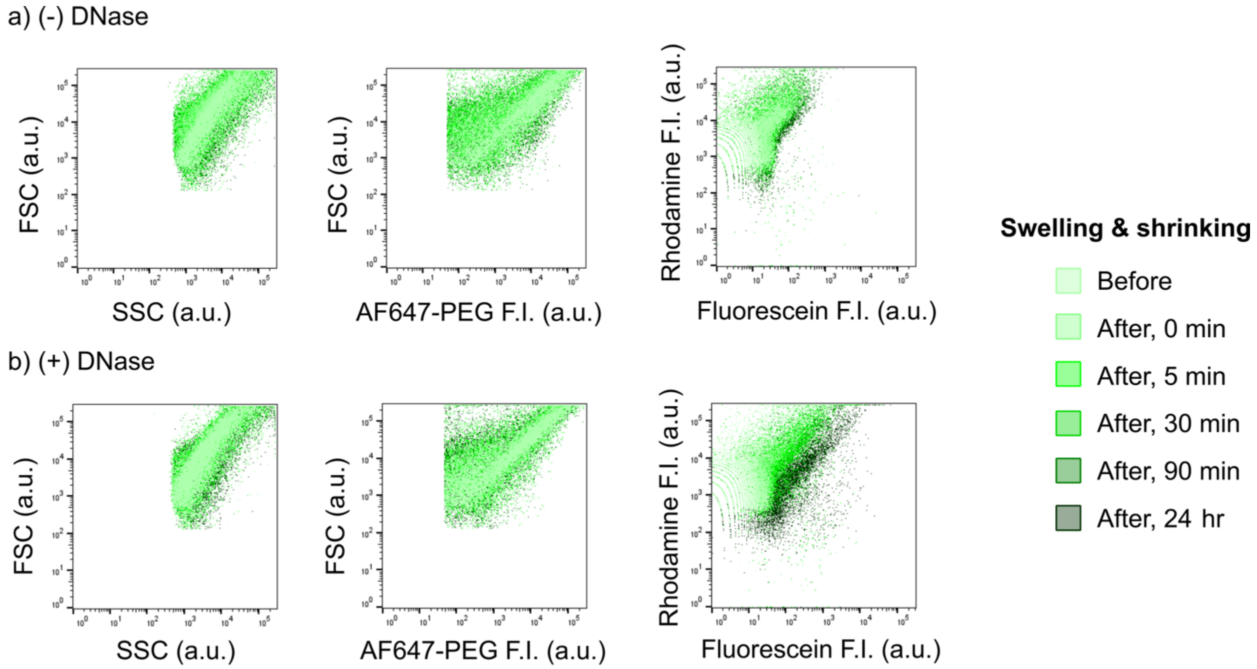


**Figure S16** - **Extended time-course data of Fig. 5d.** Two-dimensional FC dot plots for MB-DNA-fueled gel-core synthetic cells (a) without (-) or (b) with (+) encapsulation of DNase. See Main Text and Fig. 5d for details.

### Supplementary Tables

| Sample | Diffusion coefficient, $D_{app}$ (μm² s⁻¹) |
|---|---|
| Liquid core (-)mal-PE | 0.78 ± 0.07 |
| Liquid core (+)mal-PE | 0.63 ± 0.09 |
| Gel core (-)mal-PE | 0.48 ± 0.03 |
| Gel core (+)mal-PE | 0.41 ± 0.02 |

**Table S1** – Mean apparent diffusion coefficients ($D_{app}$) of NBD-PE in the membrane for each synthetic cell condition, calculated from the FRAP-derived $t_{1/2}$ values ($n$=20 for all).

| $(PEG\text{-}SH)_4$ concentration | FITC-dextran $M_w$ | Diffusion coefficient, $D_{app}$ ($\mu m^2\ s^{-1}$) |
|---|---|---|
| 1 % w/v $(PEG\text{-}SH)_4$ | 3-5 kDa | 42.24 ± 1.73 |
| | 10 kDa | 10.26 ± 0.27 |
| 5 % w/v $(PEG\text{-}SH)_4$ | 3-5 kDa | 14.97 ± 0.67 |
| | 10 kDa | 7.74 ± 0.21 |

**Table S2** – Mean apparent diffusion coefficients ($D_{app}$) of FITC-dextran in the gel-core lumen for each synthetic cell condition, calculated from the FRAP-derived $t_{1/2}$ values ($n$=20 for all).

| Fluorescent probe | $R_h$ (nm) | 2.5 % w/v PEG | 5.0 % w/v PEG | 7.5 % w/v PEG |
|---|---|---|---|---|
| Calcein | 0.7 | 0.99 ± 0.21 | 0.80 ± 0.18 | 0.83 ± 0.18 |
| 3-5 k FITC-dextran | 1.7 | 0.81 ± 0.09 | 0.72 ± 0.21 | 0.76 ± 0.14 |
| 10 k FITC-dextran | 2.7 | 0.72 ± 0.16 | 0.51 ± 0.18 | 0.51 ± 0.14 |
| 70 k FITC-dextran | 6.8 | 0.53 ± 0.08 | 0.35 ± 0.16 | 0.22 ± 0.10 |
| 500 k FITC-dextran | 17 | 0.25 ± 0.07 | 0.10 ± 0.06 | 0.12 ± 0.08 |

**Table S3** – Relative permeation $I_{in}$ / $I_{out}$ of fluorescent probes of varying size and for different PEG gel fractions. Values indicate the mean ±1 s.d.

| Fluorescent probe | $r_s / r_m$ (when $R_h \approx r_s$) | | |
|---|---|---|---|
| | **2.5 % w/v PEG** | **5.0 % w/v PEG** | **7.5 % w/v PEG** |
| Calcein | 0.04 | 0.05 | 0.05 |
| 3-5 k FITC-dextran | 0.10 | 0.13 | 0.15 |
| 10 k FITC-dextran | 0.17 | 0.20 | 0.24 |
| 70 k FITC-dextran | 0.42 | 0.53 | **0.60** |
| 500 k FITC-dextran | **1.05** | **1.32** | **1.51** |

**Table S4** – Values of solute radius ($r_s$) divided by the theoretically estimated mesh radius ($r_m$). The theoretical estimates of $r_m$ are 16.2, 12.9 and 11.2 nm respectively for [PEG-$SH_4$] at 2.5, 5.0 and 7.5 % w/v (see **Supplementary Equations S1-S2**). The Giddings model fails when $r_s / r_m > 0.6$, with these conditions highlighted in bold red in the table.

| Dilution Factor (DF) | Gel-core synthetic cell sample volume (µL) | Volume of Milli-Q added (µL) |
|---|---|---|
| 1.00 | 10 | 0.0 |
| 1.25 | 10 | 2.5 |
| 1.50 | 10 | 5.0 |
| 1.75 | 10 | 7.5 |
| 2.00 | 10 | 10.0 |
| 2.25 | 10 | 12.5 |
| 2.50 | 10 | 15.0 |
| 2.75 | 10 | 17.5 |
| 3.00 | 10 | 20.0 |

**Table S5** – Volumes used when incubating gel-core synthetic cells in increasingly hypotonic buffer (Fig. 3a-b). Table indicates the corresponding volumes used at each dilution factor (DF). Only gel-core synthetic cells encapsulating 7.5% w/v PEG-$SH_4$ were exposed to DF>2.5, since the gel-core synthetic cells at 2.5 and 5.0 % w/v PEG-$SH_4$ had already shown complete release of calcein and membrane fissure formation by DF=2.5.

| Figure | Populations compared | *p* value | Significance |
|---|---|---|---|
| 1e | liquid(-)mal-PE *vs* liquid(+)mal-PE | 0.049 | * |
| | liquid(+)mal-PE *vs* gel(-)mal-PE | 0.19 | - |
| | gel(-)mal-PE *vs* gel(+)mal-PE | 0.10 | - |
| | liquid(-)mal-PE *vs* gel(-)mal-PE | $9.2 \times 10^{-4}$ | ** |
| | liquid(+)mal-PE *vs* gel(+)mal-PE | $5.4 \times 10^{-3}$ | * |
| | liquid(-)mal-PE *vs* gel(+)mal-PE | $2.7 \times 10^{-6}$ | **** |
| 1f | 1% w/v PEG 3-5 kDa *vs*<br>10 kDa FITC-dextran | $5.7 \times 10^{-5}$ | *** |
| | 1% w/v PEG 10 kDa *vs*<br>5% w/v PEG 3-5 kDa FITC-dextran | 0.66 | - |
| | 1% w/v PEG 3-5 kDa *vs*<br>5% w/v PEG 3-5 kDa FITC-dextran | $9.8 \times 10^{-3}$ | * |
| | 5% w/v PEG 3-5 kDa *vs*<br>10 kDa FITC-dextran | 0.018 | * |
| | 1% w/v PEG 10 kDa *vs*<br>5% w/v PEG 10 kDa FITC-dextran | 0.012 | * |
| | 1% w/v PEG 3-5 kDa *vs*<br>5% w/v PEG 10 kDa FITC-dextran | $3.2 \times 10^{-5}$ | **** |
| 2c-d | **Liquid-core *vs* gel-core size changes:**<br>hypotonic | $2.5 \times 10^{-5}$ | **** |
| | **Liquid-core *vs* gel-core size changes:**<br>hypertonic | $5.1 \times 10^{-8}$ | **** |
| 3c | **Before *vs* After:** GFP influx | $3.4 \times 10^{-9}$ | **** |
| | **Before *vs* After:** GFP efflux | $1.1 \times 10^{-25}$ | **** |
| 4f | **Calcein F.I. for gel(-)mal-PE:**<br>Hypotonic *vs* Post-calcein addition | 0.016 | * |
| | **Calcein F.I. for gel(+)mal-PE:**<br>Hypotonic *vs* Post-calcein addition | $6.5 \times 10^{-3}$ | * |
| | **Calcein F.I. for post-calcein addition:**<br>gel(-)mal-PE *vs* gel(+)mal-PE | 0.029 | * |
| 5f | **(+)DNase:** 0h *vs* 24h | 0.021 | * |
| | **24h:** DNase(+) *vs* DNAse(-) | 0.040 | * |

**Table S6** – Explicit *p* values for all relevant Figures.

### Supplementary References

1. Canal, T. & Peppas, N. A. Correlation between mesh size and equilibrium degree of swelling of polymeric networks. *J. Biomed. Mater. Res.* **23**, 1183–1193 (1989).

2. Richbourg, N. R. *et al.* Precise control of synthetic hydrogel network structure via linear, independent synthesis-swelling relationships. *Sci. Adv.* **7**, 3245–3257 (2021).

### Supplementary Movies

The concentration of PEG-$SH_4$ used in all movies was 5% w/v unless otherwise specified. The synthetic cell samples were immobilised on neutravidin-coated slides by including biotinylated lipid in the synthetic cell membranes (see **Methods**).

**Movie S1: FRAP experiments using liquid-core synthetic cells**

LSCM time-lapse for a representative liquid-core synthetic cell undergoing FRAP in support of Figure 1b. The synthetic cell encapsulates rhodamine-PE in the membrane (yellow), with AF488-PEG (green) and AF647-labelled ovalbumin (red) in the lumen. The white box indicates the photobleaching ROI of AF488-PEG. Channels (L-R): rhodamine-PE (yellow), AF488-PEG (green), AF647-labelled ovalbumin (red), composite.

**Movie S2: FRAP experiments using gel-core synthetic cells**

LSCM time-lapse for a representative gel-core synthetic cell undergoing FRAP in support of Figure 1c. The synthetic cell encapsulates rhodamine-PE in the membrane (yellow), with AF488-PEG (green) and AF647-labelled ovalbumin (red) in the lumen. The white box indicates the photobleaching ROI of AF488-PEG. Channels (L-R): rhodamine-PE (yellow), AF488-PEG (green), AF647-labelled ovalbumin (red), composite.

**Movie S3, S4**: **The effect of detergent on gel- and liquid-core synthetic cells**

LSCM time-lapse in support of Figure S1, showing gel-core (S3) and liquid-core (S4) synthetic cells undergoing membrane lysis. The synthetic cells consist of rhodamine-PE in the membrane (yellow), with AF488-PEG (green) and AF647-labelled ovalbumin (OA647, red) in the lumen. The synthetic cell sample is immobilised on neutravidin-coated slides by including biotinylated lipid in the membranes. (S3) In the case of gel-core synthetic cells, when the Triton-X-100 reaches the synthetic cell sample, it lyses the lipid membranes to leave behind the AF488-labelled gel cores. Since the gel-core synthetic cells are no longer attached to the glass slide, they begin to migrate in direction of the convective flow. The OA647 is also mostly released once the membrane is lysed. (S4) In the case of liquid-core synthetic cells, the detergent lyses the lipid membranes, with complete rupture of the synthetic cells and release of the encapsulated contents.

Channels (L-R): rhodamine-PE (yellow), AF488-PEG (green), AF647-labelled ovalbumin (red), transmitted light (grey), composite.

**Movie S5: FRAP of a gel-core synthetic cell lipid membrane**

LSCM time-lapse in support of Figure 1d-e, demonstrating FRAP of NBD-PE in the membrane for a representative gel-core synthetic cell. The synthetic cell encapsulates NBD-PE in the membrane (green). FRAP is conducted as per the **Methods**, with 20 frames collected pre-bleach, the ROI then subjected to photobleaching (white box) and the recovery monitored for the remaining 60 frames. These steps are indicated in the movie. Channel: NBD-PE (green).

**Movie S6: FRAP analysis of the diffusion of molecules in gel-core synthetic cells**

LSCM time-lapse for a representative gel-core synthetic cell undergoing FRAP in support of Figure S2. The synthetic cell encapsulates AF647-labelled ovalbumin (OA647, red) and calcein (green) in the lumen. The white box indicates the photobleaching ROI of both channels. Photobleaching of both channels occurs throughout the movie, with a frame recorded after each photobleaching iteration (see **Methods**). It can be seen that the calcein signal photobleached inside the entire lumen, with the fluorescence finally completely photobleached (16 sec). This indicates that the calcein is able to freely diffuse throughout the gel core. On the other hand, the OA647 fluorescence initially reduces to a small degree both inside and outside of the photobleaching ROI, but finally photobleached just within the ROI, to leave an asymmetric fluorescence signal (20 sec). The remaining OA647 signal (red) on the left-hand side of the lumen indicates an immobilised fraction that is not photobleached - since it does not move into the photobleaching ROI. The calcein and photobleached OA647 fluorescence within the lumen reduces beyond the background fluorescence since a small amount of unencapsulated calcein and OA647 is likely present in the outer solution. Channels (L-R): AF647-labelled ovalbumin (red), calcein (green), composite.

**Movie S7: Treating gel-core synthetic cells with detergent yields membraneless hydrogels**

LSCM time-lapse in support of Figure S3, indicating that lysis of the membrane of gel-core synthetic cells leads to release of encapsulated calcein and AF647-labelled ovalbumin (OA647). The synthetic cells encapsulate rhodamine-PE in the membrane (yellow), with OA647 (red) and calcein (green) in the lumen. Triton-X-100 (2% in buffer isotonic with the synthetic cell sample) is added to the top of the imaging well. When the Triton-X-100 reaches the synthetic cell sample, it lyses the lipid membranes to leave behind the unlabelled gel cores with a portion of bound OA647 remaining (black spheres in the rhodamine channel; faint red signal in the OA647 and composite channels).

Channels (L-R): rhodamine-PE (yellow), OA647 (red), calcein (green, transmitted light (grey), composite.

**Movie S8: Shrinking of membraneless hydrogels**

LSCM time-lapse in support of Figure S3, showing the incubation of membraneless gel cores in hypertonic buffer. Gel-core synthetic cells were first prepared encapsulating AF488-PEG and AF647-labelled ovalbumin (OA647) in the lumen. These gel-core synthetic cells were loaded into an imaging well and the membranes lysed by addition of 2% Triton-X-100 to the top of the imaging well. After membrane lysis, hypertonic buffer (2x concentration of all osmolytes relative to the synthetic cell preparation buffer) was added to the top of the imaging well to give a final buffer concentration of twice the initial osmolarity. The gel cores shrink upon exposure to the hypertonic buffer, leading to an increase in the OA647 and AF488-PEG fluorescence intensities as the fluorophores are concentrated into a smaller volume. Channels (L-R): OA647 (red), AF488-PEG (green), composite.

**Movie S9: Gel-liquid transition inside synthetic cells using redox reagents**

LSCM time-lapse in support of Figure S4, of gel-core synthetic cells undergoing gel disassembly upon addition of dithiothreitol (DTT) , to yield liquid cores within the lumen. The synthetic cells encapsulate rhodamine-PE in the membrane (yellow), with AF488-PEG (green) in the lumen. The gel core is first photobleached (0-110 s) in order to demonstrate the gel state of the lumen. Membrane-permeable DTT is then added from the top of the imaging well (120 s), with gel disassembly inside the lumen evident from 220 s. Complete disassembly is achieved within 350 s, with the AF488-PEG fluorescence becoming homogenised throughout the lumen. Channels (L-R): rhodamine-PE (yellow), AF488-PEG (green), transmitted light (grey), composite.

**Movie S10a: Gel-liquid transition inside synthetic cells using redox reagents**

LSCM time-lapse in support of Figure S4, of gel-core synthetic cells undergoing gel disassembly upon addition of dithiothreitol (DTT) , to yield liquid cores within the lumen. The PEG-$SH_4$ sample is then re-oxidised as shown in Movie S10. The synthetic cells encapsulate rhodamine-PE in the membrane (yellow), with AF647-PEG (red) and calcein (green) in the lumen. The gel core (red) is first photobleached to demonstrate the gel state of the lumen. Membrane-permeable DTT is then added from the top of the imaging well, with gel disassembly inside the lumen evident as the photobleaching ROI begins to mix with the surrounding AF647-PEG. The calcein in the lumen remains encapsulated throughout, indicating that internal gel disassembly does not comprise the integrity of the lipid membrane. Channels (L-R): rhodamine-PE (yellow), AF647-PEG (red), AF488-PEG (green), transmitted light (grey).

**Movie S10: Gel-liquid-gel reversible transition inside synthetic cells**

LSCM time-lapse in support of Figure S4, indicating that the gel network inside synthetic cells can be sequentially disassembled and reassembled by redox reactions. The synthetic cells encapsulate rhodamine-PE in the membrane (yellow), with AF647-PEG (red) and calcein (green) in the lumen. The gel core is first disassembled to a liquid state using DTT (in Movie S10a). The gel network in the lumen is then reassembled by addition of $H_2O_2$ to the outer buffer, by addition from the top of the imaging well. This is done prior to the start of the movie. The return to a gel state in the lumen is then confirmed in this movie by photobleaching of the AF647-PEG in the indicated ROI (white box). Channels (L-R): rhodamine-PE (yellow), AF647-PEG (red), calcein (green), transmitted light (grey).

**Movie S11-12: Dynamics of liquid-core synthetic cells in hypotonic and hypertonic conditions**

LSCM time-lapse in support of Figure 2c, indicating the incubation of liquid-core synthetic cells in hypotonic (S11) and hypertonic (S12) conditions. The synthetic cells encapsulate rhodamine-PE in the membrane (yellow) and AF647-PEG (red) in the lumen. (S11) In the case of hypotonic conditions, the osmotic imbalance between the outer buffer and internal lumen leads to transient pore formation in the synthetic cell membranes to allow the osmolarity to re-equilibrate. This leads to some AF647-PEG release in a small fraction of the synthetic cells, but most of the synthetic cells do not show such release. (S12) In the case of hypertonic conditions, the osmotic imbalance between the outer buffer and internal lumen leads to synthetic cells deflating (20 s) and the membranes becoming floppy due to having excess surface area. The final size of the swollen synthetic cells is reduced relative to the initial size, with non-spherical membranes visible. Channels (L-R): rhodamine-PE (yellow), AF647-PEG (red), composite.

**Movie S13-14: Dynamics of gel-core synthetic cells in hypotonic conditions**

LSCM time-lapse in support of Figure 2d, indicating the incubation of gel-core synthetic cells in hypotonic conditions. The synthetic cells encapsulate rhodamine-PE in the membrane (yellow) and AF647-PEG (red) in the lumen. Milli-Q is added to the top of the imaging well to give a final buffer concentration of half the initial osmolarity. The gel cores swell in the hypotonic conditions, leading to an increase in size of the synthetic cells and slight reduction in the AF647-PEG fluorescence intensity as the fluorophore is diluted into a larger volume. Membrane fissures are visible in the rhodamine and composite channels (yellow) when the gel core swells beyond a critical volume. Channels (L-R): rhodamine-PE (yellow), AF647-PEG (red), composite.

**Movie S15-S16: Dynamics of gel-core synthetic cells in hypertonic conditions**

LSCM time-lapse in support of Figure 2d, indicating the incubation of gel-core synthetic cells in hypertonic conditions. The synthetic cells encapsulate rhodamine-PE in the membrane (yellow) and AF647-PEG (red) in the lumen. The gel cores shrink upon exposure to the hypertonic conditions, leading to shrinking of the synthetic cells and an increase in the AF647-PEG fluorescence intensity as the fluorophore is concentrated into a smaller volume. Owing to the excess membrane surface area, some tubulation of the lipid membranes is seen. Channels (L-R): rhodamine-PE (yellow), AF647-PEG (red), composite.

**Movie S17: Calcein efflux from gel-core synthetic cells**

LSCM time-lapse in support of Figure 2g, indicating the incubation of gel-core synthetic cells in hypotonic conditions, leading to membrane fissure formation and calcein release. The synthetic cells encapsulate rhodamine-PE in the membrane (yellow) and AF647-PEG (red) and calcein (green) in the lumen. Milli-Q is added to the top of the imaging well to give a final buffer concentration of half the initial osmolarity. The gel cores swell in the hypotonic buffer, leading to an increase in size of the synthetic cells. Membrane fissures are visible in the rhodamine and composite channels (yellow) when the gel core swells beyond a critical volume. Calcein efflux is seen via these membrane fissures from around 260 s onwards. Channels (L-R): rhodamine-PE (yellow), AF647-PEG (red), calcein (green), transmitted light (grey), composite.

**Movie S18: Calcein influx into gel-core synthetic cells**

LSCM time-lapse indicating the incubation of gel-core synthetic cells in hypotonic buffer containing calcein, leading to membrane fissure formation and calcein influx into the lumen. The synthetic cells encapsulate rhodamine-PE in the membrane (yellow) and AF647-PEG (red) in the lumen. Milli-Q containing calcein (10 μM) is added to the top of the imaging well to give a final buffer concentration of approximately half the initial osmolarity. The gel cores swell in the hypotonic conditions, leading to an increase in size of the synthetic cells. Membrane fissures are visible in the rhodamine and composite channels (yellow) when the gel core swells beyond a critical volume. Calcein influx is seen via these membrane fissures, causing the gel-core synthetic cell lumen to become green and hardly distinguishable against the background calcein signal. Channels (L-R): rhodamine-PE (yellow), AF647-PEG (red), calcein (green), transmitted light (grey), composite.

**Movie S19: Behavior of calcein in liquid-core synthetic cells**

LSCM time-lapse indicating the incubation of liquid-core synthetic cells containing calcein in hypotonic buffer, in support of Figure 2e. The synthetic cells encapsulate rhodamine-PE in the membrane (yellow) and AF647-PEG (red) and calcein (green) in the lumen. Milli-Q is added to the top of the imaging well to give a final buffer concentration of half the initial osmolarity. The liquid-core synthetic cells barely swell in the hypotonic buffer, leading to no efflux of calcein from the synthetic cell lumen. Channels (L-R): rhodamine-PE (yellow), AF647-PEG (red), calcein (green), transmitted light (grey), composite.

**Movie S20: Efflux of GFP from gel-core synthetic cells**

LSCM time-lapse showing the incubation of gel-core synthetic cells containing GFP in hypotonic conditions, in support of Figures 3c and S7. The synthetic cells encapsulate rhodamine-PE in the membrane (yellow) and AF647-PEG (red) and GFP (green) in the lumen. Milli-Q is added to the top of the imaging well to give a final buffer concentration of half the initial osmolarity. The gel-core synthetic cells swell in the hypotonic conditions, leading to membrane fissure formation and efflux of GFP from the lumen (around 500 s onwards). The gel-core synthetic cells appear to turn red in the composite channel as GFP is released. Hypertonic buffer (2x concentration of osmolytes used in the outer buffer for synthetic cell preparation) is then added and reaches the immobilised synthetic cells at 1120 sec, where the synthetic cells are seen to shrink, with the immobilised GFP retained. The final GFP fluorescence levels are lower than the initial levels despite the gel-core synthetic cells returning to their initial size, since much of the GFP was already released through the induced membrane fissures. Channels (L-R): rhodamine-PE (yellow), AF647-PEG (red), GFP (green), transmitted light (grey), composite.

**Movie S21: Influx of GFP into gel-core synthetic cells**

LSCM time-lapse indicating the incubation of gel-core synthetic cells in hypotonic conditions containing GFP, in support of Figures 3c and S7. The synthetic cells encapsulate rhodamine-PE in the membrane (yellow) and AF647-PEG (red) in the lumen. Milli-Q containing GFP (green) is added to the top of the imaging well to give a final buffer concentration of half the initial osmolarity. The gel-core synthetic cells swell in the hypotonic buffer, leading to membrane fissure formation and influx of GFP into the synthetic cell lumen. Channels (L-R): rhodamine-PE (yellow), AF647-PEG (red), GFP (green), composite.

**Movie S22: Behavior of GFP inside liquid-core synthetic cells**

LSCM time-lapse indicating the incubation of liquid-core synthetic cells containing GFP in hypotonic conditions, in support of Figure S9. The synthetic cells encapsulate rhodamine-PE in the membrane (yellow) and AF647-PEG (red) and GFP (green) in the lumen. Milli-Q is added to the top of the imaging well to give a final buffer concentration of half the initial osmolarity. The liquid-core synthetic cells barely swell in the hypotonic buffer, leading to no efflux of GFP from the synthetic cell lumen except for in one synthetic cell. Under subsequent exposure to hypertonic conditions (1190 s), where the synthetic cells are seen to shrink, the encapsulated GFP is retained. Channels (L-R): rhodamine-PE (yellow), AF647-PEG (red), GFP (green), transmitted light (grey), composite.

**Movie S23: Cell-free GFP synthesis in gel-core synthetic cells**

Epifluorescence microscopy time-lapse over 9.5 hrs indicating the expression of GFP in gel-core synthetic cells, in support of Figure S10. The synthetic cells encapsulate rhodamine-PE in the membrane (yellow) and AF647-PEG (red) in the lumen. The GFP fluorescence (green) is seen to increase over time as the protein is expressed. Channels (L-R, top-to-bottom): DIA (grey), GFP (green), rhodamine-PE (yellow), AF647-PEG (red).

**Movie S24: Cell-free GFP synthesis in liquid-core synthetic cells**

Epifluorescence microscopy time-lapse over 9.5 hrs indicating the expression of GFP in liquid-core synthetic cells, in support of Figure S10. The synthetic cells encapsulate rhodamine-PE in the membrane (yellow) and AF647-PEG (red) in the lumen. The GFP fluorescence (green) is seen to increase over time as the protein is expressed. Channels (L-R, top-to-bottom): DIA (grey), GFP (green), rhodamine-PE (yellow), AF647-PEG (red).

**Movie S25: Calcein exchange between gel-core synthetic cells and the environment**

LSCM time-lapse indicating the release and capture of calcein from gel-core synthetic cells via osmotic swelling and shrinking, in support of Figure 4e. The synthetic cell encapsulates rhodamine-PE in the membrane (yellow) and calcein (green) in the lumen. Milli-Q is added to the top of the imaging well to give a final buffer concentration of half the initial osmolarity. The gel-core synthetic cell swells in the hypotonic buffer, leading to membrane fissure formation and efflux of calcein into the external buffer (0-120 s). Addition of hypertonic buffer of twice the initial osmolarity is then added, leading to the gel-core synthetic cells shrinking and the membrane fissures resealing. Calcein in isotonic buffer is then added, leading to no gel-core synthetic cell swelling or shrinking and it is seen that the calcein does not permeate the gel core, indicating that the membrane fissures have resealed (130-450 s). Hypotonic buffer containing is then added again, causing the gel-core synthetic cell to swell and calcein influx (460-680 s). Subsequent shrinking in hypertonic buffer leads to membrane resealing with calcein retainment in the lumen (690-1690 s). Final washing with isotonic buffer causes no change in the synthetic cell size, and washing away of external calcein (1700 s onwards). Channels (L-R): rhodamine-PE (yellow), calcein (green), composite.

**Movie S26: β-gal activity in gel-core synthetic cells containing mal-PE**

LSCM time-lapse in support of Figure 5b, indicating the uptake of FDG into gel-core synthetic cells containing immobilised β-galactosidase (β-gal) in the gel network and mal-PE in the membrane. Gel-core synthetic cells were first prepared encapsulating AF647-PEG and β-gal in the lumen (see **Methods**). These gel-core synthetic cells were loaded into an imaging well and hypotonic buffer (0.5x concentration of all osmolytes relative to the synthetic cell preparation buffer) containing the non-fluorescent molecule FDG (0.75 mM) added to the top of the imaging well to give a final buffer concentration of half the initial osmolarity. The gel cores swell upon exposure to the hypotonic conditions, leading to membrane fissure formation and the influx of FDG into the gel-core synthetic cell lumen. Addition of hypertonic buffer (2x concentration of all osmolytes relative to the synthetic cell preparation buffer) restores the initial osmolarity and induces membrane resealing. Over time, the up-taken FDG is hydrolysed to fluorescein by the immobilised β-gal, leading to an increase in the green fluorescence intensity. Channels (L-R): rhodamine-PE (yellow), AF647-PEG (red), fluorescein (green), transmitted light (grey), composite.

**Movie S27-28: β-gal activity in liquid and gel-core synthetic cells**

LSCM time-lapse in support of Figure 5b, showing (S27) the uptake of FDG into gel-core synthetic cells containing immobilised β-galactosidase (β-gal) in the gel network but without mal-PE in the membrane and (S28) the limited uptake of FDG into liquid-core synthetic cells containing β-galactosidase (β-gal) in the lumen and mal-PE in the membrane. Synthetic cells were first prepared encapsulating AF647-PEG and β-gal in the lumen. FDG was added as described in **Movie S28**. Over time, there is limited change in the fluorescein fluorescence of the synthetic cells. However, up-taken FDG is hydrolysed to fluorescein by the co-encapsulated β-gal in a minor population in both S29 and S30 (leading to an increase in the green fluorescence intensity). Channels (L-R): rhodamine-PE (yellow), AF647-PEG (red), fluorescein (green), transmitted light (grey), composite.
